# Interferon-independent STING enforces epithelial genome-integrity checkpoint to restrain tumor evolution

**DOI:** 10.64898/2026.02.17.706442

**Authors:** Hang Xiang, Martina Macino, Clemens Woitaske-Proske, Nicholas Bodenstein, Joana P Bernardes, Florian Tran, Nicole A. Schmid, Gabriela Rios Martini, Felix Wottawa, Johanna Bornhaeuser, Nassim Kakavand, Oliver Dyck Dionisi, Silke van den Bossche, Guang Yang, Qicong Wu, Julia Kugler, Lina Welz, Bjorn Konukiewitz, Julius C. Fischer, Christian Peifer, Stefan Schreiber, Philip Rosenstiel, Markus Tschurtschenthaler, Ashley D Sanders, Konrad Aden

## Abstract

STING is canonically known for mediating interferon responses to cytosolic DNA, yet its cell-intrinsic role in genome maintenance beyond the immune context is unknown. Here we show that epithelial STING functions as a type I interferon-independent genome-integrity checkpoint. STING loss impairs homologous recombination repair, attenuates ATM-associated damage signaling, elevates CDK1 activity, and causes chromosomal instability revealed by single-cell Strand-seq, culminating in spontaneous intestinal adenocarcinoma. These defects arise before tumor formation and confer selective vulnerability to CDK inhibition in tumor organoids and human colorectal cancer cells. Our findings identify STING as a cell-autonomous guardian of epithelial genome stability that restrains chromosomal instability-driven tumor evolution beyond its canonical immune function.

## Main Text

The intestinal mucosa is a highly proliferative tissue that constantly renews the epithelial lining into specified cell types to meet the different physiological demands in a healthy state (*1*). However, during DNA replication, DNA polymerases frequently misincorporate ribonucleotides into the genome, and in the absence of efficient ribonucleotide excision repair (RER) these lesions can accumulate to >1,000,000 sites per genome, making them one of the most abundant naturally occurring replication errors (*2*). Failure of RER, which removes misincorporated ribonucleotides from genomic DNA via the enzyme RNase H2, leads to DNA damage and subsequent activation of p53 as a major tumor suppressor, resulting in loss of proliferation and induction of cellular senescence (*2, 3*). We and others have shown that several key cellular homeostatic factors (ER-stress, Autophagy) are able to modify the p53-dependent tumor suppression, resulting in genomic instability and transcription-associated mutagenesis (*4, 5*). Importantly, failure of RER not only leads to induction of DNA damage confined to the nucleus but also activates the cGAS-STING pathway through to the accumulation of cytosolic micronuclei containing damaged dsDNA (*6*). dsDNA sensing by cGAS-STING pathway results in the activation of a canonical innate immune response, involving phosphorylation of Tank-binding protein (TBK), Interferon-related factor 3 (IRF) and subsequent induction of type I interferon (IFN-I) (*7*). Beyond its canonical role in innate immune signaling, cGAS itself has been described to modulate DNA repair pathways. In both mice and humans, cGAS can inhibit homologous recombination (HR) by interfering with RAD51-mediated repair, whereas a 4 amino acid change in the naked mole cGAS potentiates DNA repair via enhanced homologous recombination, which may contribute to its cancer resistance (*8, 9*). Notably, the reported cGAS-STING effects on HR appear to be context-dependent and remain controversial. Along the same line, cGAS-derived cGAMP has been reported to trigger DNA damage signaling, supporting a broader role of the cGAS-STING axis in genome maintenance (*10*). However, the exact mechanism by which STING-dependent dsDNA sensing facilitates DNA repair is not understood. In addition, the anti-tumor role of cGAS-STING is mostly viewed in the context of anti-tumor immunity in immune cells, in which therapeutic stimulation of STING is investigated as an adjunct therapy in cancers such as prostate cancer (*11–14*). In addition, whether STING also acts in an epithelial-intrinsic manner to sense DNA damage induced by RER failure and preserve genome integrity is not known. It has been previously reported that functional STING signaling is abrogated in colorectal cancer, but whether this is mechanistically linked to preservation of genome integrity is unknown (*15, 16*). Here, we identify epithelial STING as a cell-intrinsic genome-stability factor that supports ataxia-telangiectasia mutated (ATM) activation and HR competence, restrains chromosomal instability, and suppresses intestinal tumorigenesis.

### Epithelial STING protects against spontaneous intestinal tumorigenesis

It has been previously established that RNase H2b deficiency induces the accumulation of cytosolic dsDNA, which potentially activate the cGAS-STING pathway (*17*). We first assessed epithelial STING expression in the intestine of 8-12-week-old *H2b*^ΔIEC^ mice, in which *Rnaseh2b* is selectively deleted in intestinal epithelial cells, and compared with *H2b*^fl/fl^ control mice (*3, 4*). *H2b*^ΔIEC^ mice showed a significant increase in Crypt-based intestinal epithelial cells expressing STING, compared to *H2b*^fl/fl^ controls (**Fig. 1A, B**), along with elevated STING protein levels in small intestinal organoids derived from *H2b*^ΔIEC^ mice (**Fig. 1C**). These findings suggest a direct association between epithelial DNA damage and STING induction.

**Fig. 1.**
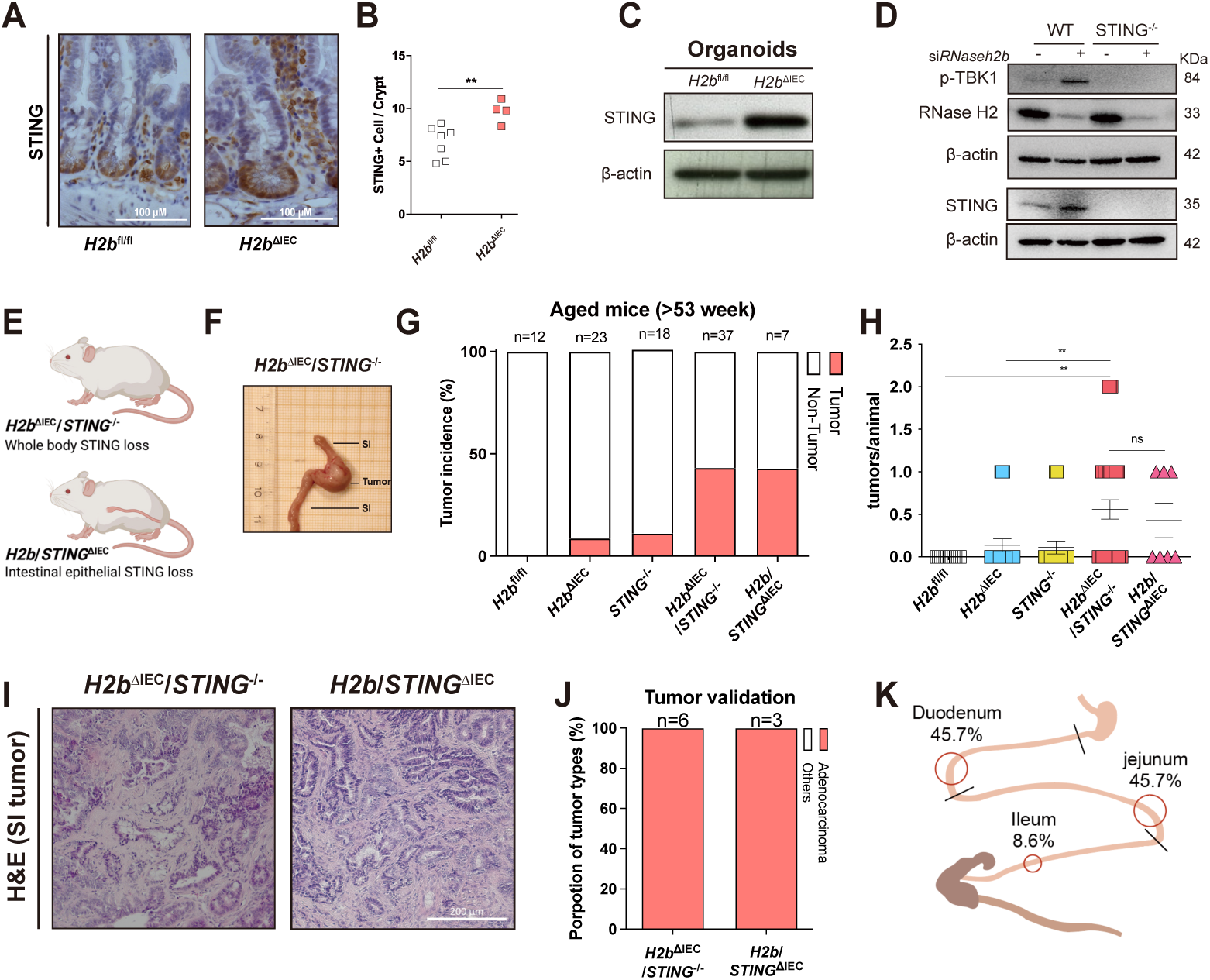
Loss of STING promotes intestinal tumorigenesis and progression in the context of epithelial DNA damage. **(A** and **B)** Immunohistochemical staining (**A**) and statistical analysis (**B**) of STING staining in the small intestine of 8-12 week old *H2b*^fl/fl^, (n=7), *H2b*^ΔIEC^ (n=4) mice. The average number of STING-positive cells per crypt was determined by counting ≥30 crypts per sample, and the mean number of positive cells per crypt was calculated. Significance was determined using unpaired two-tailed Student’s test. **(C)** Western Blot analysis of STING in small intestinal organoids from 8-12 week old *H2b*^fl/fl^ and *H2b*^ΔIEC^ mice. **(D)** MODE-K WT and STING^-/-^ cells were transfected with either siRNA against *Rnaseh2b* or control siRNA for 24h. Protein levels were measured by western blot assay. **(E)** Schematic of generation of *H2b*^ΔIEC^/*STING*^-/-^ mouse line. **(F)** Representative picture of small intestinal tumor from aged (>53W) *H2b*^ΔIEC^/*STING*^-/-^ mouse. **(G** and **H)** Tumor incidence (G) and tumors per mouse (H) in *H2b*^fl/fl^ (n=12), *H2b*^ΔIEC^ (n=23), *STING*^-/-^ (n=18), *H2b*^ΔIEC^/*STING*^-/-^ (n=37), and *H2b*/*STING*^ΔIEC^ (n=7) mice. Significance was determined using unpaired two-tailed Student’s test. **(I)** H&E staining of *H2b*^ΔIEC^/STING^-/-^ and *H2b*/*STING*^ΔIEC^ tumor **(J)** Histopathological tumor grading confirms presence of adenocarcinoma of *H2b*^ΔIEC^/STING^-/-^ and *H2b*/*STING*^ΔIEC^ tumor. **(K)** Scheme of the mouse small intestine showing regional tumor incidence based on the corresponding quantification data from *H2b*^ΔIEC^/STING^-/-^ mouse. For all the significance analysis: ns = not significant, * p<0.05, ** p<0.01, *** p<0.001, **** p<0.0001.

To investigate a potential link between DNA damage repair and cGAS-STING signaling in intestinal epithelial cells (IECs), we generated CRISPR/Cas9-mediated MODE-K knockout lines targeting either *Tmem173* (STING^-/-^) or cGAS (cGAS^-/-^) (**fig. S1A-B**). DNA damage was induced by either *Rnaseh2b* knockdown or cytarabine (AraC) treatment. Under both conditions, cGAS^-/-^ and STING^-/-^ cells showed markedly reduced cGAS-STING signaling, including diminished TBK1 phosphorylation and reduced induction of interferon inducible genes (**Fig. 1D, fig. S2A-G**). These findings indicate that RER impairment activates cGAS-STING-dependent signaling in intestinal epithelial cells.

To investigate the tumor-suppressive role of epithelial STING *in vivo* in the context of RNase H2b deficiency-induced DNA damage, we generated *H2b*^ΔIEC^/*STING*^-/-^ mice in which Tmem173 is additionally ablated and monitored intestinal tumor development in aged animals (>53 weeks) (**Fig. 1E, fig. S1C-D**). *H2b*^ΔIEC^/*STING*^-/-^ mice exhibited a markedly increased tumor incidence (16/37) compared with *H2b*^fl/fl^ (0/12), *H2b*^ΔIEC^ (2/23), or *STING*^-/-^ (2/18) controls, along with a significant increase of tumors per mouse (**Fig. 1F-H**). Notably, all intestinal tumors in *H2b*^ΔIEC^/*STING*^-/-^ mice were adenocarcinomas localized to the small intestine, predominantly affecting the duodenum (45.7%) and jejunum (45.7%) (**Fig. 1I-K**). In addition, *H2b*^ΔIEC^/*STING*^-/-^mice exhibited shortened colon and small intestine length, while body and liver weight were unchanged compared to *H2b*^fl/fl^ (**fig. S3A-D**).

To test whether STING loss within the intestinal epithelium is sufficient to drive tumorigenesis under DNA damage, we generated mice with epithelial-specific deletion of STING on the *H2b*^ΔIEC^ background (*H2b*/*STING*^ΔIEC^). At baseline, these mice closely resembled systemic knockout animals (**fig. S3A-H**) and developed spontaneous small-intestinal adenocarcinomas (3/7) with comparable incidence and histology (**Fig. 1G-J**), indicating that tumorigenesis arises from epithelial STING loss.

As additional evidence of aggressiveness, tumor-derived organoids from *H2b*^ΔIEC^/STING^-/-^ mice formed malignant orthotopic tumors after transplantation into the colonic mucosa of immunodeficient NSG mice, and organoids from one primary tumor gave rise to liver metastases in recipient animals (**fig. S3I-L**). Collectively, these findings identify STING as a protective factor in the intestinal epithelium that limits chronic DNA damage-driven transformation and restrains spontaneous small-bowel adenocarcinoma formation.

### Loss of STING results in chromosomal instability and clonal expansion in intestinal epithelium

The spontaneous tumor phenotype prompted us to investigate whether STING loss destabilizes the epithelial genome at single-cell resolution. To directly assess this, we performed Strand-seq, to quantify the rate and formation of genetic aberrations on a cell-by-cell basis in small intestinal organoids derived from *H2b*^fl/fl^*, H2b*^ΔIEC^ and *H2b*^ΔIEC^/*STING*^-/-^ mice (**fig. S4A**). Strand-seq is a single-cell DNA sequencing method designed to detect structural rearrangements and sister chromatid exchanges (SCEs) arising in mitotic cells (*18–21*). Because SCEs reflect crossover events generated during HR repair of double-strand break (DSB), and manifest as strand-state changes or breakpoints (BPs) in Strand-seq data (*21*), they provide a quantitative readout of HR-associate genome instability (**fig. S4B-C, fig. S5**).

We found a significant increase in the total number of BPs per cell in *H2b*^ΔIEC^ cells (median= 6, range= 1-113, p= 7.6e-07) as compared to *H2b*^fl/fl^ control cells (median = 3, range= 0-10) indicating an elevated rate of SCE formation under *H2b* deficiency. This phenotype was further exacerbated by additional STING loss with *H2b*^ΔIEC^/*STING*^-/-^ cells showing a median= 44 BPs per cell (range= 5-67, p= <2.22e-16) (**Fig. 2A-B**). To clearly distinguish genomically unstable cells, we defined a threshold of >10 BPs based on the upper distribution of *H2b*^fl/fl^ cells. Using this cutoff, *H2b*^ΔIEC^ cells segregated into two subpopulations, with 34% classified as unstable (>10 BPs; median = 41 BPs/cell; p= 3.72e-10 vs *H2b*^fl/fl^), whereas 90% of *H2b*^ΔIEC^/*STING*^-/-^ cells exceeded this threshold (median = 45 BPs/cell) (**Fig. 2B-C**). Notably, among the unstable cells, *H2b*^ΔIEC^ and *H2b*^ΔIEC^/*STING*^-/-^ groups exhibited comparable SCE burdens (p= 0.61) (**Fig. 3D**). Indicating that STING loss does not increase the instability per cell but instead expands the proportion of cells that acquire elevated genomic instability.

**Fig. 2.**
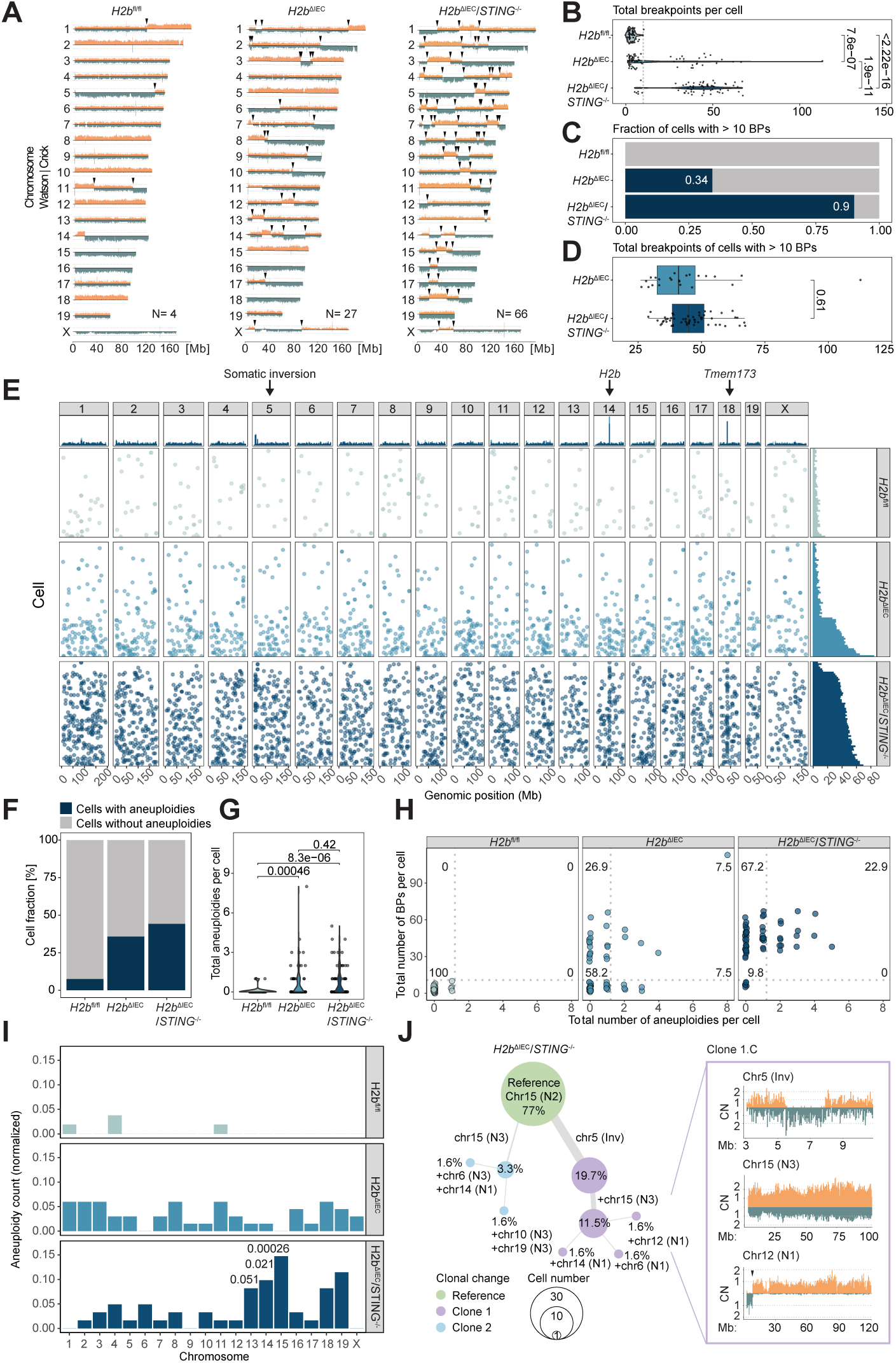
Loss of STING results in chromosomal instability and clonal expansion in intestinal epithelium. **(A)** Representative ideograms of single cells Strand-seq libraries for *H2b*^fl/fl^, *H2b*^ΔIEC^ and *H2b*^ΔIEC^/*STING*^-/-^. The ideograms of each chromosome are showing binned read counts aligned to the mouse reference genome mm09. The colors indicate the positive (Crick) in blue and the negative (Watson, orange) template strand. The black arrows are marking somatic changes in directionalities, or breakpoints. **(B** to **D)** Quantification of breakpoints and genomic instability. (B) Violin plot showing the manual quantifications of somatic breakpoints per cell by each condition. Each dot represents one single cell. (C) Stack plot showing the fraction of cells with more than 10 BPs in blue and less or equal to 10 BPs in grey. (D) Boxplot of the total number of breakpoints per cell of cells with more than 10 BPs. The P-value was calculated using the two-sample t-test. **(E)** The plot is showing the breakpoints per cell by genomic location, identified with BreakpointR and filtered for recurrent regions identified in the *H2b*^fl/fl^ cells. Each dot represents a breakpoint. On the top panel the stack-histogram shows the quantification per genomic position (500Kb bins). The panel on the right shows the histogram of the quantification of total breakpoint per cell. Chromosome Y was removed due to the low read mappability. **(F)** Histogram showing the percentage of cells that possess at least one aneuploidy. **(G)** Violin plot showing the number of aneuploidies per cell. Each dot represents one single cell. The P-value was calculated with the two-sample t-test. **(H)** Dot plot of the correlation between the count of breakpoint per cell and aneuploidy per cell. The dotted grey lines mark the upper range found in *H2b*^fl/fl^ cells (BPs= 10, Aneuploidies= 1). The numbers shown are the percentage of cells in each quadrant. **(I)** Histogram of total numbers of aneuploidies per chromosome normalized by the cell number per condition. The P-value was calculated on the single-cell counts using one-way analysis of variance (ANOVA) and pairwise comparisons using pairwise t-tests with Benjamini-Hochberg (BH) correction for *H2b*^ΔIEC^/*STING*^-/-^ vs *H2b*^ΔIEC^. **(J)** Clonal expansion in *H2b*^ΔIEC^/*STING*^-/-^ cells. Right panel: Network plot showing the clonal fraction of cells with alterations. The black arrow is marking a somatic breakpoint. Left panel: representative ideograms of regions affected in Clone 1.C.

**Fig. 3.**
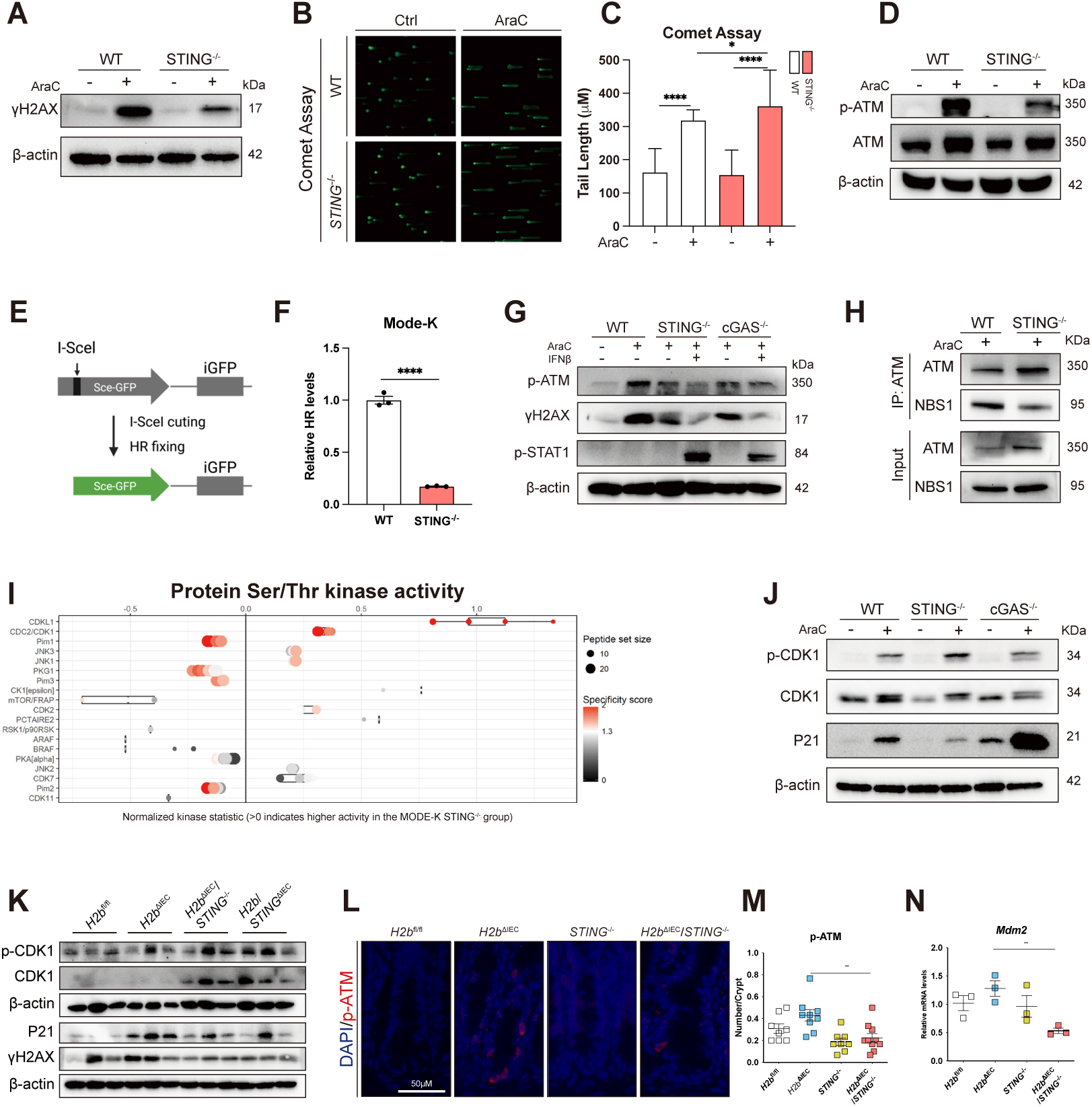
STING sustains homologous recombination and G2/M checkpoint control independent of IFN-I. **(A)** Western blot analysis of γH2AX from MODE-K WT and STING^-/-^ cell lines after treated with 2.5 μM AraC for 20h. **(B** and **C)** Representative picture (B) and statistical analysis of tail length (C) measured from comet assay from MODE-K WT and STING^-/-^ cell lines after treated with 2.5 μM AraC for 20h. Significance was determined using unpaired two-tailed Student’s test. **(D)** Western blot analysis of ATM from MODE-K WT and STING^-/-^ cell lines after treated with 2.5 μM AraC for 20h. **(E** and **F)** Scheme of DR-GFP reporter assay (E). HR levels of MODE-K WT and STING^-/-^ cell lines were measured by FACS (F). Significance was determined using unpaired two-tailed Student’s test. **(G)** MODE-K WT and STING^-/-^ cells were transfected with either siRNA against *Wip1* or control siRNA for 24h then stimulated with 2.5 μM AraC for 20h. Protein levels were measured by western blot assay. **(H)** Co-immunoprecipitation (co-IP) analysis of the interaction between ATM and NBS1 from MODE-K WT and STING-/- cell lines after treated with 2.5 μM AraC for 20h. ATM-containing complexes were immunoprecipitated using an anti-ATM antibody and probed for NBS1 by immunoblotting. **(I)** Deeper look into the results from upstream kinase analysis showing the main affected STK, the full figure shown in **fig. S9D**. **(J)** MODE-K WT, STING^-/-^, and cGAS^-/-^ cells were treated with 2.5 μM AraC for 20h. Protein levels were analyzed by western blot. **(K)** Protein levels from young (8-12W) *H2b*^fl/fl^, *H2b*^ΔIEC^, *H2b*^ΔIEC^/*STING*^-/-^, *H2b*/*STING*^ΔIEC^ mouse small intestine crypt sample assessed by western blot assay. **(L** and **M)** Immunohistochemistry and statistical analysis of the small intestine of aged (>53W) H2bfl/fl (n=8), *H2b*^ΔIEC^ (n=9), STING^-/-^ (n=8) and *H2b*^ΔIEC^/*STING*^-/-^ (n=10) mice, stained for p-ATM. Representative images (L) of p-ATM immunofluorescence staining (red) and corresponding statistics (M). The average number of p-ATM-positive cells per crypt was determined by counting ≥30 crypts per sample, and the mean number of positive cells per crypt was calculated. Significance was determined using unpaired two-tailed Student’s test. **(N)** Gene expression of *Mdm2* in *H2b*^fl/fl^, *H2b*^ΔIEC^, *STING*^-/-^ and *H2b*^ΔIEC^/*STING*^-/-^ mice assessed by qPCR taqman probe assay, n=3. Significance was determined using unpaired two-tailed Student’s test. For all the significance analysis: ns = not significant, * p<0.05, ** p<0.01, *** p<0.001, **** p<0.0001.

To assess whether SCEs preferentially occurred at specific genomic locations, such as fragile sites or recombination hotspots, we mapped BPs genome-wide using *Breakpoint.R* (*22*) and removed germline-associated BPs, defined as recurrent regions (>3 BPs per bin) detected in control cells (**Fig. 2E, fig. S3A-B**). Despite the significantly elevated SCE burden in *H2b*^ΔIEC^ and *H2b*^ΔIEC^/*STING*^-/-^ cells, BPs were broadly distributed across the genome, indicating stochastic, genome-wide instability rather than localized fragility. Three notable enrichments were observed: two distinct peaks at the *H2b* locus on Chr14 (*H2b*^ΔIEC^ and *H2b*^ΔIEC^/*STING*^-/-^) and at the *Sting (Tmem173)* locus on Chr18 (*H2b*^ΔIEC^/*STING*^-/-^ only), likely reflecting rearrangements introduced during targeted gene deletion (**Fig. 3E, fig. S3, fig. S4**); and a third enriched region that corresponded to a subclonal inversion on Chr5 (Chr5:4,988,964-5,923,558). This inversion is unique to *H2b*^ΔIEC^/*STING*^-/-^ cells and expanded to a cell fraction (CF) of 0.197 (n=12) (**Fig. 2E, fig. S4**), consistent with positive selection.

We next assessed whether STING loss induces structural chromosomal abnormalities, including aneuploidies, associated with oncogenic transformation. Both *H2b*^ΔIEC^ and *H2b*^ΔIEC^/*STING*^-/-^ cells showed a marked increase in aneuploidy frequency compared to *H2b*^fl/fl^ controls: 7.55% of *H2b*^fl/fl^ cells possess aneuploidies, versus 35.8% in *H2b*^ΔIEC^ and 44.3% in *H2b*^ΔIEC^/*STING*^-/-^ cells, corresponding to a 4.7-fold (p= 9.04e-05 ) and a 5.9-fold (p= 2.98e-06) increase, respectively (**Fig. 2F**). However, the difference between the two knockout populations was modest, and the overall number of aneuploidies per cell did not differ significantly between *H2b*^ΔIEC^/*STING*^-/-^ and *H2b*^ΔIEC^ cells (p = 0.42) (**Fig. 2F-G**). These data indicate that STING loss does not substantially elevate whole-chromosome copy number changes beyond those induced by *H2b* deficiency alone. Integrating SCE burden with aneuploidy frequency, revealed a pronounced shift in the *H2b*^ΔIEC^/*STING*^-/-^ population. Whereas the majority of *H2b*^ΔIEC^ cells remained genomically stable (58.2% in the low-BP/low-aneuploidy quadrant), *H2b*^ΔIEC^/*STING*^-/-^ cells exhibited extensive genomic instability (67.2% in the high-BP/low-aneuploidy quadrant), with a marked increase in cells harbouring both high SCE levels and aneuploidies (22.9% vs 7.5% in *H2b*^ΔIEC^) (**Fig. 2H**).

Having learned that aneuploidies accumulate in both *H2b*^ΔIEC^ and *H2b*^ΔIEC^/*STING*^-/-^cells with only minor differences in overall frequency (1.2-fold, p= 0.335), we next investigated whether specific chromosomes were disproportionately affected. *H2b*^ΔIEC^/*STING*^-/-^ cells displayed a significant enrichment of aneuploidies affecting chromosomes 14 (p= 0.021) and 15 (p= 0.00026) compared with *H2b*^ΔIEC^ cells (**Fig. 2I**). Chromosome 15 is of particular interest as it harbors key cancer-associated genes such as *Myc* and *Rad21* and has been previously linked to tumorigenesis in mouse models (*23*). Trisomy of Chr15 was detected in 14.8% of *H2b*^ΔIEC^/*STING*^-/-^ cells and arose independently in two distinct lineages (**Fig. 2J**). In “clone 1”, a Chr5 inversion (present in 19.7% of cells, as described above) preceded the gain of Chr15 (CF = 0.12), followed by additional chromosomal alterations involving chromosomes 6, 12, and 14 (CF= 0.016, each). In “clone 2”, Chr15 gain occurred independently (CF= 0.33), and was followed by further alterations on chromosomes 6, 10, 14, and 19. The recurrent and independent formation of Chr15 trisomies, together with their subsequent clonal expansion and acquisition of additional aneuploidies in *H2b*^ΔIEC^/*STING*^-/-^ cells, suggests that loss of both *H2b* and *Sting* not only promotes aneuploidy but also creates a permissive and selective environment for clonal evolution and ongoing chromosomal alterations.

### STING deficiency impairs homologous recombination in intestinal epithelial cells

The extensive chromosomal instability observed upon STING loss suggested a failure of genome maintenance mechanisms. We therefore examined the extent to which STING directly regulates DNA damage response (DDR) in IECs. Following induction of DNA damage with AraC treatment, MODE-K WT cells exhibited a marked increased γH2AX signal, but not in STING^-/-^ cells (**Fig. 3A**), a finding that was further confirmed by immunofluorescence staining (**fig. S8A-B**). Because γH2AX primarily reflects DNA damage signaling and repair rather than DNA break formation per se (*24*), we next assessed the amount of DNA breaks by performing comet assays. STING^-/-^ cells showed even higher levels of DNA damage after AraC treatment, compared to MODE-K WT (**Fig. 3B-C**). Taken together, these results indicate that despite increased DSB formation in STING^-/-^ cells, STING deficiency results in impaired γH2AX induction.

Following DSB formation, HR repair is initiated by ATM dimer activation through auto-phosphorylation at Ser1981, enabling phosphorylation of downstream DDR targets such as H2AX (*25, 26*). To determine whether the reduced γH2AX signal reflects impaired ATM activation, we assessed ATM phosphorylation and observed markedly reduced p-ATM levels in MODE-K STING^-/-^ cells compared to WT following AraC treatment (**Fig. 3D**). To functionally validate defective HR repair, we employed the DR-GFP reporter assay, which quantifies HR efficiency by measuring restoration of functional GFP after I-Scel-endonuclease induced DSBs (*27*) (**Fig. 3E**). Consistent with impaired ATM activation, STING deficiency resulted in significantly reduced HR efficiency (**Fig. 3F**).

To determine whether this defective DDR requires the entire cGAS-STING pathway, we repeated these experiments in MODE-K cGAS^-/-^ cells. Similar to STING loss, cGAS deficiency resulted in γH2AX induction (**fig. S8B-C**) and diminished ATM phosphorylation (**fig. S8D**), despite comparable levels of DNA damage (**fig. S8E**). These findings indicate that an intact cGAS-STING axis is required for efficient DNA damage signaling in IECs.

We next investigated whether IFN-I, the canonical downstream effector of cGAS-STING signaling (*28, 29*), mediates this DDR function. Notably, exogenous IFN-β treatment failed to rescue γH2AX induction or ATM phosphorylation (**Fig. 3G**), and knockdown of *Ifnar1* has no effect on DDR protein levels (**fig. S8F**). These results indicate that cGAS-STING coordinates DDR independently of IFN-I signaling.

To further define the downstream pathway involved, we performed siRNA-mediated knockdown of *Tbk1* and *Irf3*. *Tbk1* knockdown reduced ATM phosphorylation but did not affect γH2AX induction, whereas *Irf3* knockdown had no detectable effect on either parameter (**fig. S8G-H**), suggesting that cGAS-STING signaling contributes to DDR initiation up to the level of TBK1 but independently of IRF3-mediated transcription.

We next explored the mechanism underlying ATM activation in MODE-K STING^-/-^cells. We first assessed WIP1 (PPM1D), a DDR phosphatase that promotes cell cycle checkpoint termination by dephosphorylating multiple ATM pathway components (*30*). Upon AraC treatment, WIP1 levels decreased in MODE-K WT cells but were increased in both cGAS^-/-^ and STING^-/-^ cells, correlating with reduced p-ATM levels (**fig. S8I**). However, siRNA-mediated knockdown of *Wip1* in STING^-/-^ cells only partially restored γH2AX levels and failed to rescue ATM phosphorylation, indicating that aberrant WIP1 activity is not the primary cause of defective ATM activation in the absence of STING (**fig. S8J**) (*30*).

Finally, we asked whether impaired ATM phosphorylation reflects defective ATM recruitment to DNA damage sites. During HR, ATM is recruited to DSBs by the MRN (MRE11-RAD50-NBS1) complex, with NBS1 providing a critical C-terminal ATM-binding interface required for efficient ATM docking and activation (*31–33*). Consistent with this model, co-immunoprecipitation experiments revealed a markedly reduced interaction between NBS1 and ATM in MODE-K STING^-/-^ cells compared with WT controls (**Fig. 3H**).

Collectively, these data demonstrate that STING loss compromises ATM-dependent DDR initiation by weakening NBS1-mediated ATM recruitment, while simultaneously altering DDR signal termination.

### STING deficiency disrupts the G2/M checkpoint response in IECs

To identify the regulatory network underlying the STING-dependent DDR, we used MODE-K WT and STING^-/-^ cells and performed kinase activity profiling using a PamGene assay to quantify phosphorylation dynamics of tyrosine kinases (PTKs) and serine/threonine kinases (STKs) kinases. STING loss was associated with a global reduction in both PTK and STK kinase activity under basal (untreated) conditions (**fig. S9A, B**). Following AraC-induced DNA damage, PTK activity was also globally reduced, whereas STK responses were heterogeneous, showing both increases and decreases (**fig. S9A-C**). Consistent with impaired IFN-I signaling, JAK1 and JAK2 activities were diminished. Similar trends were also found for several DNA damage-associated STKs, including IKKα, IKKβ, and ATR. In contrast, kinases linked to cell-cycle progression and proliferation, including CDK1, CDK7, and CDKL1, were upregulated in AraC-treated STING^-/-^ cells. Among these, Cyclin-Dependent kinase 1 (CDK1) exhibited the strongest increase in kinase activity, based on kinase activity and phosphorylation status prediction (**Fig. 3I, fig. S9D-F**). Pathway enrichment analysis using EnrichR further revealed a significant downregulation of the p53 pathway in AraC-treated STING^-/-^ cells (**fig. S9G**). The p53-p21 axis is a central regulator of the G2/M DNA damage checkpoint, acting to restrain CDK1 activity and prevent premature mitotic entry, whereas CDK1, a key Ser/Thr kinase, drives G2-to-M transition once checkpoint constraints are released (*34, 35*). To determine whether STING deficiency alters this checkpoint balance, we measured CDK1 and p21 levels in MODE-K cells after DNA damage. Consistent with the kinase screening results, STING^-/-^ cells displayed elevated CDK1 phosphorylation and reduced p21 levels confirming the attenuated p53 target genes such as *Mdm2* or *Ddit4l* (**Fig. 3J, fig. S10A-B**). These data indicate that loss of STING impairs the p53-p21 arm of the DNA damage response and shifts cells toward G2/M checkpoint failure. Notably, although cGAS deficiency also impaired ATM activation, it did not reproduce the downstream p21/CDK1 imbalance observed in STING-deficient cells, suggesting that this checkpoint defect is selectively linked to STING (**Fig. 3J, fig. S10C-D**).

To determine whether these defects arise as an early, cell-intrinsic consequence of STING loss rather than secondary effects of tumor progression or aging, we examined DDR and G2/M checkpoint markers in intestinal crypts from young *H2b*^ΔIEC^/*STING*^-/-^and *H2b*/*STING*^ΔIEC^ mice prior to detectable tumor formation. Consistent with our observations in MODE-K cells, both *H2b*^ΔIEC^/*STING*^-/-^ mice and intestinal epithelium-specific *H2b*/*STING*^ΔIEC^ mice displayed elevated CDK1 levels together with reduced p21 and γH2AX in intestinal crypts compared with *H2b*^ΔIEC^ controls (**Fig. 3K**), indicating a cell-autonomous defect in checkpoint control in vivo. Importantly, these alterations persisted in aged *H2b*^ΔIEC^/*STING*^-/-^ mice, which exhibited reduced ATM phosphorylation and attenuated p53 signaling, further supporting impaired DDR checkpoint enforcement upon epithelial STING loss (**Fig. 3L-N**).

### STING deficiency defines a chromosomal-instability-prone epithelial state vulnerable to CDK inhibition

Given the increased CDK1 activity and impaired genome maintenance observed above, we asked whether STING-deficient tumors adopt a transcriptional program indicative of CDK-driven proliferation. RNA sequencing of intestinal organoids derived from *H2b*^ΔIEC^ and *H2b*^ΔIEC^/*STING*^-/-^ mice revealed distinct global transcriptomic profiles between the two genotypes (**Fig. 4A-C**).

**Fig. 4.**
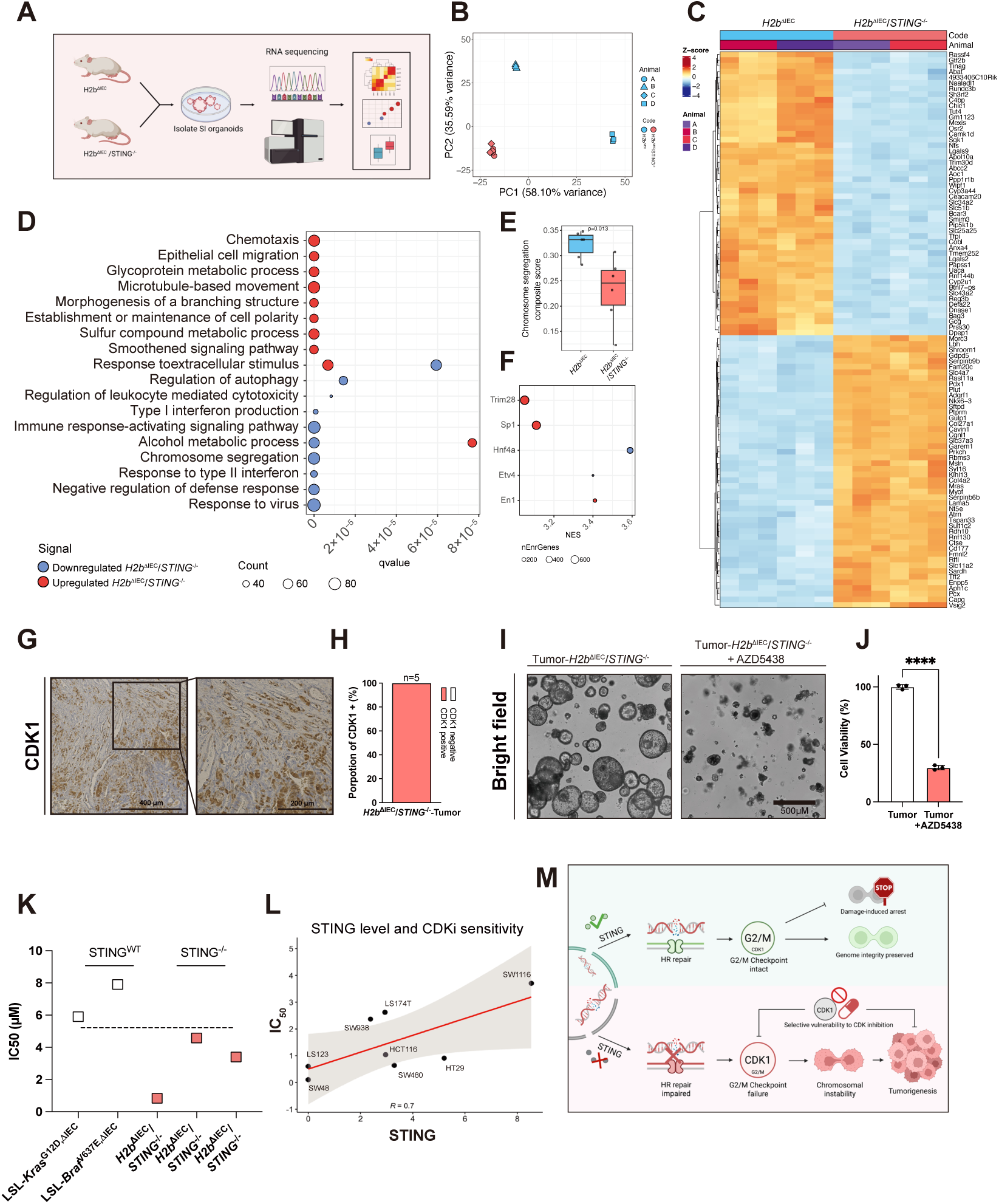
STING loss rewires DNA damage–associated transcriptional programs and confers vulnerability to CDK inhibition. **(A** to **F)** RNA-seq identifies distinct transcriptional programs in *H2b*^ΔIEC^ and *H2b*^ΔIEC^/*STING*^-/-^ Organoids. Scheme of RNA-seq model (A). Principal component analysis (PCA) of RNAseq samples. *H2b*^ΔIEC^/*STING*^-/-^ samples clustered tightly and were independent of the animal of origin, whereas *H2b*^ΔIEC^ samples segregated from *H2b*^ΔIEC^/*STING*^-/-^ along both principal components and also separated by animal of origin along PC2 (B). Heatmap of the top 50 differentially expressed genes (DEGs) between *H2b*^ΔIEC^ and *H2b*^ΔIEC^/*STING*^-/-^ group. DEG expression profiles were highly concordant between animals within each genotype, with robust differences between genotypes; in *H2b*^ΔIEC^/*STING*^-/-^ group, 2,388 genes were upregulated and 2,612 genes were downregulated relative to *H2b*^ΔIEC^ (C). Gene Ontology (GO) overrepresentation analysis for DEGs upregulated (red) and downregulated (blue) in *H2b*^ΔIEC^/*STING*^-/-^group. Dot size indicates the number of genes associated with each enriched term. The top 10 GO terms per direction were selected based on term dispensability calculated using rrvgo (D). Transcription factor binding site (TFBS) enrichment analysis for DEGs upregulated (red) and downregulated (blue) in *H2b*^ΔIEC^/*STING*^-/-^ group. Dot size indicates the number of genes associated with each enriched transcription factor (E). Single-sample GSEA (ssGSEA) score for nuclear chromosome segregation. Based on the RNA-seq data described above, ssGSEA was performed to calculate pathway activity scores for nuclear chromosome segregation using the MSigDB gene set GOBP_NUCLEAR_CHROMOSOME_SEGREGATION (F). **(G** and **H)** Immunohistochemical staining of CDK1 in tumor tissues derived from H2b^ΔIEC^/STING^-/-^ mice. Representative picture of CDK1 staining(G). Proportion of CDK1-positive tumors (H). **(I** and **J)** Representative pictures (I) and CellTiter-Glo assay (J) from *H2b*^ΔIEC^/*STING*^-/-^ tumor organoids treated with Control (DMSO) or 1 μM AZD5438 for 72h. Significance was determined using unpaired two-tailed Student’s test. **(K)** Comparison of AZD5438 IC_50_ values in STING^WT^ and STING^-/-^ tumor organoids. IC_50_ values for AZD5438 were determined from dose–response assays and compared between STING^WT^ tumor organoids and STING^-/-^ tumor organoids based on the dose-response curve shown in **fig. S12B** to **F**. **(L)** Correlation between AZD5438 sensitivity and STING expression in CRC cell lines. Linear regression analysis was performed to assess the relationship between AZD5438 sensitivity (IC_50_ values in different colorectal cancer cell lines; **fig. S12G** to **N**) and STING protein levels determined by western blot (**fig. S12O**; quantified by densitometry as indicated). **(M)** Proposed model for epithelial STING function in genome integrity control. In epithelial cells, STING supports HR repair and maintains G2/M checkpoint integrity, thereby enabling damage-**induced** arrest and preserving genome stability. Loss of STING impairs HR repair, weakens G2/M checkpoint control in association with increased CDK1 activity, and promotes chromosomal instability, ultimately contributing to tumorigenesis. This checkpoint-defective state is accompanied by selective vulnerability to CDK inhibition. For all the significance analysis: ns = not significant, * p<0.05, ** p<0.01, *** p<0.001, **** p<0.0001.

In addition to the downregulation of interferon signaling and host immune defense pathways, gene ontology analysis revealed a marked reduction in chromosome segregation and mitotic spindle organization signatures in *H2b*^ΔIEC^/*STING*^-/-^ organoids (**Fig. 4D**). To quantify this finding, we calculated a chromosome segregation signature score and found it markedly reduced in *H2b*^ΔIEC^/*STING*^-/-^ organoids (**Fig. 4E**), suggesting disrupted mitotic fidelity, which may underlie the observed aneuploidy accumulation.

Transcription factor activity analysis further revealed substantial rewiring of regulatory networks in *H2b*^ΔIEC^/*STING*^-/-^ organoids. Regulons associated with TRIM28, SP1, and EN1 were enriched, whereas those linked to HNF4A and ETV4 were reduced (**Fig. 4F**), indicating altered transcriptional control of epithelial identity and cell-cycle-related processes. Collectively, these changes suggest a transcriptional state compatible with dysregulated proliferation and genome maintenance.

These transcriptional changes suggested that STING-deficient tumors may rely on CDK-driven mitotic progression. We therefore tested the effects of pharmacological CDK inhibition. Firstly, we examined CDK1 expression in tumors and found all tumor samples derived from both *H2b*^ΔIEC^/*STING*^-/-^ mouse (n=5) and *H2b*/*STING*^ΔIEC^ mouse (n=3) were positive for CDK1 (**Fig. 4G-H, fig. S11A-B**). Subsequently, organoids established from these tumors exhibited complete loss of proliferative capacity upon treatment with the pan-CDK inhibitor AZD5438 (**Fig. 4I-J, fig. S11C-D**). Based on the STING-CDK1 signature, we asked whether STING expression governs the sensitivity of tumor cells towards CDK inhibitor. In MODE-K cells, STING^-/-^ cells were significantly more sensitive than WT or cGAS^-/-^ controls (**fig. S12A**). We further compared the IC_50_ values in murine tumor organoids from *H2b*^ΔIEC^/*STING*^-/-^ mice with murine tumor organoids harboring *Kras^G12D^* or *Braf^V637E^*mutations and confirmed STING-genotype specific sensitivity to CDK inhibition (**Fig. 4K, fig. S12B-F**). Extending these findings to human disease, we tested AZD5438 across multiple human adenocarcinoma cell lines in which STING deficiency has been systematically assessed previously. We observed that the sensitivity was inversely correlated with STING expression levels, indicating that reduced STING expression predicts enhanced responsiveness to CDK inhibition (*16*) (**Fig. 4L, fig. S12G-O**). Collectively, these results identify defective STING-mediated DDR as a key driver of hyperproliferation and chromosomal instability and uncover CDK inhibition as a potential therapeutic vulnerability in STING-deficient tumors.

## Discussion

While the mechanistic concept of DNA damage, cytosolic dsDNA accumulation and cGAS-STING dependent sensing is widely established across several species and cell types, the immediate role of the cGAS-STING pathway on preserving genome integrity has remained inconclusive (*36–39*). Here, we challenge the prevailing view of STING as primarily an immunostimulatory sensor and reposition STING as an epithelial, IFN-I-independent regulator upstream of HR.

Using *H2b*^ΔIEC^/*STING*^-/-^ mice model, we show that these mice develop spontaneous intestinal adenocarcinoma and further transform into metastatic tumors. Mechanistically, we show that cGAS/STING deficiency reduces the interaction between ATM and NBS1, a central component of the MRN complex, thereby attenuating ATM recruitment and activation. This finding underscores a critical role for the cGAS/STING axis in initiating DSB-induced DDR. In parallel, loss of cGAS/STING enhanced γH2AX dephosphorylation through upregulation of the phosphatase WIP1, further compromising DNA damage signaling. Together, these findings reveal a dual function of cGAS-STING in coordinating both the activation and maintenance of DDR.

To explicitly resolve the consequence of defective STING on genome integrity, we employed Strand-seq and RNA-seq. Our data revealed that double deficiency of H2b and STING may induce extensive sister chromatid exchanges and chromosome-specific aneuploidies, e.g. trisomy 15, which harbors oncogenic drivers such as *Myc* and *Rad21*. These genomic alterations coincide with downregulation of chromosome segregation and mitotic spindle genes, and reprogramming of transcription factors including TRIM28, SP1, and EN1, indicating a shift toward a dedifferentiated, DNA damage-tolerant state. Importantly, this transcriptional reprogramming was accompanied by elevated CDK activity and impaired G2/M checkpoint control, consistent with a hyperproliferative phenotype. Together, these findings delineate a mechanistic cascade whereby defective DNA repair and cell-cycle control synergize to promote chromosomal instability, clonal selection and tumor evolution.

We wondered whether the very same properties that promote STING-dependent tumor formation may also generate a mechanistic vulnerability that can be targeted therapeutically. We could demonstrate that STING-deficiency results in increased CDK1 activation. Importantly, we observed a striking dichotomy between STING and cGAS, as cGAS deficient cells retained functional p53 signaling.

Notably, CDK inhibition effectively suppresses the growth of H2b/STING-deficient organoids and tumors, establishing aberrant CDK signaling as a critical vulnerability in this context. Extending these findings to human cancer models, STING-low adenocarcinoma cell lines displayed heightened sensitivity to CDK inhibition, suggesting that defective STING-mediated DNA damage response dictates both proliferative drive and therapeutic susceptibility. Our findings have fundamental implications. First, while pan-CDK inhibitors have been hindered by dose-limiting toxicity (*40*), our data suggest that the intrinsic CDK dependency of STING-deficient tumors transforms this toxicity challenge into a dosage-selectivity opportunity (*15, 41*). Potential limitations might be the fact that single-strand seq was conducted in *H2b*^ΔIEC^/*STING*^-/-^ but not in *H2b*/*STING*^ΔIEC^. However, we point out that tumor manifestation incidence and CDK1 activation was comparable in both genetic models, arguing for an epithelial-cell intrinsic function of STING.

Our results not only provide a mechanistic rationale for targeting CDK pathways in STING-deficient intestinal tumors but also highlight a broader principle in which loss of genome maintenance reprograms cell-cycle control to fuel oncogenic adaptation.

## Materials and Methods

### Cell culture

Mouse intestine epithelial cell line MODE-K (WT and Knock out) was cultured in DMEM GlutaMAX-I (Gibco, #61965-026) medium containing 10% fetal bovine serum (Sigma, #F7524), 1% HEPES 1M (Gibco, #15630-056), 1% NEAA 100x (Gibco, #11140-035) and 1% penicillin streptomycin (Gibco, #15140-122) at 37℃ in a constant temperature incubator with 5% CO2.

### Generating of MODE-K cGAS^-/-^ and STING^-/-^ cell line by using Crispr-Cas9 system

Single-stranded DNA with guiding sequence target for mice *cGAS* (Fw 5′-AGATCCGCGTAGAAGGACGAGTTTT-3′; Rev 3′-TCGTCCTTCTACGCGGATCTCGGTG-5′) and *Tmem173* gene (Fw 5’-CAGTAGTCCAAGTTCGTGCGGTTTT-3’, Rev 5’-CGCACGAACTTGGACTACTGCGGTG-3’) were annealed and cloned into CRISPR nuclease vector via GeneArt® CRISPR Nuclease Vector with OFP Reporter Kit (Thermofisher, A21174). Vector with target sequence were then transformed into TOP 10 competent *E. coli* (Thermofisher, C404003) and selected via ampicillin LB solid medium. The selected bacteria were then enriched and proceeded with plasmid extraction by using HiSpeed Plasmid Midi Kit (Qiagen, 12643) for further transfection. MODE-K WT cells were seeded one day before transfection, and transfected purified plasmid DNA into the cells on the next day by using Lipofectamine^TM^ 3000 Reagent (Thermofisher, L3000015), and incubated in 37℃, 5% CO2 incubator overnight. After 24h of incubation, cells transfected with CD4 reporter vector were enriched and selected by CD4 Dynabeads. Selected cells then diluted and seeded into 96 well plates with 1 cell each well. After cultured for 7-14 days, cells with single colony were selected out, those cells were enrichment cultured and further tested by Western Blot.

### Immunofluorescence for cells

Cells were seeded onto autoclaved coverslips within a cell culture plate. Following the desired stimulation or transfection, the cells were fixed and permeabilized using -20°C methanol for 10 minutes at room temperature. After washing with PBS, the cells were blocked using 2% BSA solution for 1 hour. Subsequently, the cells were subjected to overnight incubation at 4°C with the primary antibody (**table S1**). The next day, after washing away unbound primary antibody, the cells were incubated with the appropriate secondary antibody (**table S1**) for 1 hour at room temperature while protecting them from light. Additional washing steps with PBS were performed to remove excess secondary antibody. To visualize the nuclei, the cells were stained with a fluorescent nuclear dye such as DAPI or Draq5, and then a final wash with PBS was carried out. Finally, the coverslips were mounted onto microscope slides for microscopy.

### Gene expression Taqman assay

Total RNA was isolated from all biopsies and samples (cells, organoids, and mouse tissues) using the RNeasy Kit (Qiagen, #74106) according to the manufacturer’s instructions. First-strand cDNA was synthesized using the Maxima H Minus First Strand cDNA Synthesis Kit (Thermo Fisher Scientific, #K1652) following the manufacturer’s protocol. Quantitative PCR was performed using TaqMan Gene Expression Assays (**table S2**). Reactions (10 µL total volume) contained 5 µL cDNA (2 ng/µL) and 5 µL TaqMan Master Mix supplemented with 0.5 µL TaqMan assay primer/probe and 4.5 µL mix, and were run on a Roche real-time PCR system (LightCycler® 96 Instrument, Roche). Relative gene expression was calculated using *β**-actin*** (*Actb*) as the endogenous reference gene.

### Western blot

Cell lysates were extracted and prepared in RIPA buffer (Thermofisher, #89901) containing 1% Phosphatase Inhibitor Cocktail (Sigma, # P5726). The concentration of protein was measured by lowry assay (Biorad, #5000116). Equal amounts of proteins (15-20μg) were loaded and separated by SDS-PAGE gel, then transferred onto PVDF membrane (BIO-RAD, # 162-0177). The membranes were blocked by 5% non-fat milk (Santa-Cruz, # sc-2325) in TBST for 1h and incubated with specific primary antibody (**table S1**) overnight in 4℃ or 1h in room temperature. Subsequently, the membranes were incubated with HRP-coupled secondary antibody (**table S1**) for 1h in room temperature and developed by ChemiDocTM imaging system (Biorad, #12003263) with ECL reagent (Biorad, #1705062).

### Cell IC50 analysis

IC50 values for AZD5438 were determined using CellTiter-Glo-based viability assay (Promega, #G9042) in 96-well format. Cells were seeded at 1.2 × 10^4 cells per well in white 96-well plates in 100 µL complete medium. After 24 h, the medium was replaced with fresh medium containing a serial dilution of AZD5438, and cells were incubated for an additional 72 h. CellTiter-Glo reagent (100 µL) was then added directly to each well, followed by incubation for 20 min at room temperature. For cell cultured in transparent plate, remove the medium and add 100 µL CellTiter-Glo reagent and transfer to a white 96-well plate after 20min incubation. Luminescence was measured using a microplate reader. Dose-response curves and IC50 values were calculated in GraphPad Prism using nonlinear regression.

### Co-Immunoprecipitation

Cells were lysed in NP-40 lysis buffer containing 1% protease inhibitor on ice for 30 min, followed by centrifugation at 12,000 × g for 10 min at 4 °C. Protein concentrations were determined by standard assays. Equal amounts (200-500 ng) of total protein were incubated with primary antibody (1:100) **(table S1)** or species-matched IgG as a negative control overnight at 4 °C. Magnetic beads (30 µL per sample) were washed three times with NP-40 lysis buffer and blocked with 2% BSA for 30 min on ice, then added to the protein-antibody mixtures and incubated for 4h at 4 °C. Beads were collected using a magnetic rack, washed three times with NP-40 lysis buffer, and proteins were eluted in 30 µL of 1× SDS loading buffer by heating at 95 °C for 5 min. 10% of total lysate was used as input. Samples were analyzed by SDS-PAGE followed by western blotting.

### Comet assay

MODE-K cells were seeded in 6-well plates and treated with either PBS (vehicle control) or 2.5 µM cytarabine (AraC) (Sigma-Aldrich, C6645) for 20 h. Cells were then harvested, and viable cell numbers were determined using an automated cell counter (LUNA II, Logos Biosystems) prior to comet assay analysis. DNA damage was quantified using an alkaline comet assay (Comet Assay® Kit, 4250-050-ESK; R&D Systems) according to the manufacturer’s instructions. Briefly, low-melting-point agarose (LMA) was melted at 90 °C and equilibrated to 37 °C. Cells were mixed with LMA, and 100 µL of the suspension was applied to CometSlides and spread evenly. Agarose gels were allowed to solidify at 4 °C for 10 min. Slides were then incubated in lysis buffer at 4 °C overnight. Following lysis, DNA was unwound in alkaline unwinding solution for 20 min. Electrophoresis was performed in alkaline running buffer at 21 V for 30 min at 4 °C. Slides were washed twice with deionized water and once with 70% ethanol, then stained with SYBR Gold (Invitrogen, S1194) for 10 min in the dark. After rinsing with deionized water and drying at 37 °C, comets were imaged using an ECHO microscope and comet tail length was quantified.

### Protein kinase assay

MODE-K WT and STING^-/-^ cells after treated with control (PBS) or 2.5 µM AraC (Siggma-Aldrich, C6645) for 20h were lysed based on the manufacturers protocol (PamGene Protocol 1160) with 100 µL MPer (Thermo Scientific) spiked with 1:100 Halt® Phosphatase-Inhibitor-Cocktail (Thermo Scientific) and 1:100 EDTA-free Halt® Protease-Inhibitor-Cocktail (Thermo Scientific). Protein concentrations of the samples were evaluated with Lowry assay.

Kinase activity profiling was performed with Tyrosine Kinase and Serine-Threonine Kinase PamChip-4® Arrays (PamGene) using a PamStation® 12. Each chip has four arrays and the arraýs surface consists of porous silica with 196 (PTK) or 144 (STK) distinct peptide targets, composed of 13 amino acids mimicking phosphorylation sites, that are printed in spots on the porous silica surface. For the tyrosine kinase analysis, a FITC-conjugated antibody was used to quantify the phosphorylation level. Furthermore, the kinetic of the reaction was measured by pumping the sample fluid through the porous surface for 94 cycles. From cycle 37 to 92 every five cycles and at cycle 94 five pictures each with different exposure times (10 ms, 20 ms, 50 ms, 100 ms and 200 ms) were taken. For the serine-threonine kinase analysis, a primary antibody was used and after cycle 90 a secondary FITC-antibody against the primary antibody was given on the arrays for quantification. From cycle 92 to 122 every five cycles and after cycle 124 five pictures each with different exposure times (10 ms, 20 ms, 50 ms, 100 ms and 200 ms) were taken. The brightness of each spot was measured against reference spots and used for further analysis. The kinase activity profiling was performed using 2 biological replicates distributed each on different chips and arrays with 5 µg (PTK) or 1 µg (STK) protein per array and a final concentration of 400 µM ATP according to the manufacturers protocol. BioNavigator® 6 (PamGene) was used for signal evaluation and upstream kinase analysis following the manufacturers protocol. Data from the upstream kinase analysis was integrated into EnrichR. Threshold cut-off for the median finale score was made at 1,2 or 1,0, respectively, and median kinase statistic results arranged from up to down before converting the UniPROT IDs from the main kinases with lower activity into gene names using UniProt. Gene names were pasted into EnrichR to draw a connection between the genes and their corresponding pathways.

### DR-GFP HR reporter assay

MODE-K WT and STING^-/-^ cells were seeded into 24-well plates in triplicate and cultured in serum-free medium. Cells were transiently transfected with the DR-GFP reporter plasmid (Addgene, #26475) using LipofectamineTM 3000 Reagent (Thermofisher, L3000015) according to the manufacturer’s instructions. After 24 h, cells were re-transfected with DR-GFP together with the I-SceI expression plasmid (Addgene, #26477) to induce site-specific DSBs. 48 hours after the second transfection, cells were harvested, and GFP-positive cells were quantified by flow cytometry (SONY FACS SA3800) DSB repair efficiency was calculated based on the percentage of GFP-positive cells as described previously.

### Generating of knockout mice

Epithelial deficiency of *Rnaseh2b* (*H2b*^ΔIEC^) mice were generated as described previously (*3*). Mouse with full-body deficiency of *Tmem173* (*STING*^-/-^) were generated by Crispr/Cas9 system while *STING*^fl/fl^ mice were obtained from the Jackson Laboratory (#:031670) to generate intestinal epithelial specific deficiency of *Tmem173*. *STING*^-/-^ and *STING*^ΔIEC^ mice were paired with *H2b*^ΔIEC^ mice to generate *H2b*^ΔIEC^/*STING*^-/-^ and *H2b*/*STING*^ΔIEC^ respectably. All mice were housed in a specific pathogen-free (SPF) animal facility at Kiel University under standard conditions, with ad libitum access to food and water and a 12 h light/dark cycle. Age-matched female and male mice were used for all experiments. Animal experiments with basal phenotype were performed in accordance guidelines with Kiel University (animal vote IX 552-241316/2022).

### Tissue processing

Mouse small intestine and colon were cut and opened longitudinally, then followed by Swiss rolling from the distal to proximal part. All the mouse tissue was fixed by 10%(w/v) formalin for 24h at 4°C. Fixed tissues were embedded in paraffin blocks and were cut for H&E and IHC staining according to the method described before (*42*).

### H&E stanning

Paraffin-embedded tissue sections were dewaxed in xylene substitute (Roti-Histol; Carl Roth) for 10 min, rehydrated through graded ethanol (3 × 1 min in 100%, 2 × 1 min in 96%, 1 × 1 min in 70%), and washed in distilled water for 5 min with one solution change on a shaker. Sections were stained with Mayer’s hematoxylin solution (Hämalaun sauer nach Meyer; Carl Roth) for 3 min (without shaking), rinsed under running tap water for 5 min for bluing, and counterstained with 0.1% aqueous eosin G (prepared by dilution from 1% stock; Carl Roth) for 3 min, followed by a 30 s rinse under running tap water. Slides were dehydrated in ascending ethanol (10 dips in 70%, 3 × 10 dips in 96%, 2 × 10 dips in 100%, then 3 × 2 min in 100%), cleared in xylene substitute (10 dips in the first bath, then 2 × 2 min in fresh baths), and mounted with Roti-Histokitt (Carl Roth); mounted slides were left to evaporate/dry for at least 24 h before further handling. Tumor classification on H&E-stained sections was performed by a pathologist according to the latest WHO Classification of Tumors of the Digestive System (*43*).

### DAB staining

Paraffin-embedded sections were dewaxed in xylene substitute (Roti-Histol; Carl Roth, #6640.2) for 10 min, rehydrated through graded ethanol (3 × 60 s in 100%, 1 × 60 s in 96%, 1 × 60 s in 70%), and rinsed briefly in distilled water, followed by PBS washes (3 × 4 min). Antigen retrieval was performed in citrate buffer (10 mM citric acid, pH 6) by heating in a pressure cooker for 20 min, then sections were washed again in PBS (3 × 4 min). Endogenous peroxidase activity was quenched with 3% H₂O₂ for 10 min, followed by PBS washes (3 × 4 min). Sections were blocked for 30 min with 1% BSA in PBS containing 0.2% Triton X-100, excess solution was removed, and primary antibodies (**table S1**) diluted in 1% BSA/PBS were incubated overnight at 4°C in a humidified chamber. After PBS washes (3 × 4 min), sections were incubated with biotinylated secondary antibodies (**table S1**) for 45 min, washed in PBS (3 × 4 min), and incubated with VECTASTAIN Elite ABC reagent for 45 min. After PBS washes (3 × 4 min), signal was developed using a DAB peroxidase substrate solution (Vector SK-4100; prepared in 5 mL distilled water with buffer, DAB, and peroxidase) until the desired staining intensity was reached, then rinsed in distilled water (5 min). Sections were counterstained with Mayer’s hematoxylin for 1 min and blued under running tap water for 5 min, dehydrated in ascending ethanol (70%, 96%, 100%), cleared in xylene substitute, and coverslipped.

### Immunofluorescence for mouse tissue

Deparaffinization, rehydration, and heat-induced antigen retrieval were performed as described for the DAB staining protocol above. After antigen retrieval, sections were then blocked for 30 min in 5% BSA in PBS containing 0.2% Triton X-100, incubated with primary antibodies (**table S1**) diluted in 1% BSA/PBS overnight at 4°C, and subsequently incubated with Alexa Fluor-conjugated secondary antibodies (1:500, 45 min, dark) (**table S1**). Nuclei were counterstained with DAPI (1:40,000 in PBS, 10 min, dark), followed by PBS washes, and slides were mounted with fluorescence mounting medium and stored at 4°C. The prepared slides were proceeded to microscopy and images were acquired using a Zeiss AxioImager.Z1 microscope equipped with Apotome and AxioVision Rel. 4.9 software (ZEISS, Oberkochen, Germany).

### Organoid Isolation

Mice were euthanatized according to animal care of the Christian-Albrechts-University (Kiel, Germany), and separate ileum out from small intestine. Remove fat as much as possible and washed by cold PBS. Cut the ileum longitudinally and cut them into 5mm small pieces after washed by cold PBS to remove the mucus and feces. Incubate these ileum spices in 5mM ice cold EDTA for 10min then shaking intermittently, the discard the supernatant which containing villus and repeat this step for 3 times. Discard supernatant and replaced by cold PBS then shake for 30s and incubate for 5min, repeat this step for 3 times. Screen the supernatant through a 100nm cell strainer and spin down with 400g for 5min in 4℃. Resuspend the pellet with Advanced DMEM F12, after counted by microscope, seed the crypts into cell culture plate with Matrigel and cultured with conditional hEGF-Noggin-Rspondin (ENR) medium.

### Organoid Passage

Discard the old medium and remove the Matrigel containing organoids with cold PBS into a 15ml falcon, then pipette it up and down to break organoids. Spin down by and discard the supernatant, and seed organoids with Matrigel to a cell culture plate, then place the plate into 37℃ an incubator for 20 to 40min. After the Matrigel became hard, add ENR medium.

### Organoid IC50 analysis

Organoid viability and IC50 were determined in tumor-derived organoids. Organoids were dissociated with TrypLE Express (Gibco, Thermo Fisher Scientific) into single cells or small clusters, resuspended in Matrigel, and seeded in triplicate using pooled material from 3-4 confluent wells of a 24-well plate for six drug concentrations. Seeding density was adjusted visually under brightfield microscopy to ensure comparable density across conditions, and Matrigel was mixed thoroughly during dispensing to minimize uneven distribution. After seeding, organoids were cultured overnight in ENR medium supplemented with Y-27632 (abcam, ab120129). On the following day (Day 0), organoids were treated with increasing concentrations of AZD5438 (0.008, 0.04, 0.2, 1, 5, and 20 μM) or vehicle control. After 72 h of treatment, cell viability was assessed using the CellTiter-Glo assay according to the manufacturer’s instructions. Luminescence was measured following transfer to a white 96-well plate, and normalized viability values were used to calculate IC50 using GraphPad Prism.

### Organoid colony forming assay

Organoid colony-forming assay was performed by dissociating organoids into single cells using TrypLE Express (Gibco, Thermo Fisher Scientific). A total of 10,000 cells were embedded in Matrigel and cultured in ENR medium supplemented with Y-27632 (abcam, ab120129), with medium changed every other day. Stimulants were added on Day 2 as indicated. On Day 3, medium was replaced with standard ENR without Y-27632. Colony formation was assessed on Day 7 by colony counting or by measuring viability using the CellTiter-Glo assay.

### Orthotopic organoid transplantation

Orthotopic transplantation was performed using two independent organoid lines derived from two individual mice and carried out according to previous description (*4*).

Each organoid line was transplanted into four recipient immunocompromised NSG (NOD.Cg-Prkdcscid Il2rgtm1Wjl/SzJ, Stock No: 005557) mice which were purchased from Charles River Laboratories (Research Models and Services, Germany GmbH, Sulzfeld, Baden-Württemberg, Germany) and housed, bred and maintained in SPF facilities (ZPF, TUM University Hospital, Technical University of Munich, Germany). Orthotopic organoid transplantation was performed as previously described (*44, 45*). Briefly, prior to transplantation, organoids were dissociated to generate small clusters (approximately 5-10 cells) using TrypLE Express (Gibco, Thermo Fisher Scientific) and suspended in a reduced-volume injection medium composed of Advanced DMEM/F12 supplemented with B27 (1×), N2 (1×), L-glutamine (Gibco, Thermo Fisher Scientific), 10% (v/v) Matrigel (Corning), 1% (v/v) penicillin/streptomycin, and 10 μM Y-27632 (STEMCELL Technologies). For each injection, 50 organoid clusters were prepared in a total volume of 100 μL while each mouse was administered 2-3 injections. Recipient mice were anesthetized, and the colonic lumen was gently flushed with PBS using a syringe fitted with a straight gavage needle. Endoscopy-guided delivery was performed with a rigid mini-endoscope system (Karl STORZ; 1.9 mm diameter; ColoView). Organoid suspensions were deposited into the colonic submucosa using a flexible 33G needle (Hamilton; custom 16-inch length; 45° bevel). Orthotopic tumor formation and metastatic lesions were subsequently assessed, then proceed with H&E staining and microscopy analysis. Data were analyzed using GraphPad Prism. The animal studies were conducted in compliance with European guidelines for the care and use of laboratory animals and were institutionally approved by the Institutional Animal Care and Use Committees (IACUC) of Technical University of Munich and by the District Government of Upper Bavaria.

### Strand Sequencing (Strand-seq)

Strand-seq library preparation: Small intestine organoids were isolated from an aged mouse (>53W, n=1) and cultured using ENR medium as described above. After passaging (P1), organoids were left to grow for 4 days before Bromodeoxyuridine (BrdU; Sigma, B9285; 40 μM) was added to the culture media for 8, 12 or 16h (as shown in fig. S7). The organoids were then dissociated into single cells using TrepLE Express (#12604021) and stored at -80°C in freezing media containing 10% DMSO and 90% FBS. To sort BrdU+ cells, the samples were thawed at 37°C, and washed once by adding 4 ml of 0.01% BSA (Thermofisher,15260037) in 1X PBS (Sigma, 882114) and centrifuged at 500g for 5 minutes. The supernatant was then decanted and the cells were resuspended in 200μl of nuclei staining buffer (100 mM Tris-HCl pH7.4, 154 mM NaCl, 1 mM CaCl2, 0.5 mM MgCl2, 0.2% BSA, 0.1% NP-40, ultra pure water, 10 ug/ml Hoechst 33258, 10 ug/ml PI) as described in previous publications (*22*). Single nuclei from the appropriate time point of BrdU (fig. S9) were sorted using the cell sorter BD FACSAria Fusion, in 96-well plates containing 5μl of freshly prepared Freeze buffer (ProFreeze_CDM (2X), 100% DMSO, 1X PBS). The plates were stored at -80°C until library preparation was performed, as described previously (*22*).

Libraries were sequenced on an Illumina NovaSeq X Plus sequencing platform using pair-end sequencing protocol (75-bp), demultiplexed and aligned to the mouse reference genome (mm09).

Strand-seq analyses, breakpoint and aneuploidies counts: High quality libraries, obtained from cells that incorporated BrdU for one complete round of DNA replication, were selected as described previously (*22*). Sharp changes in read alignment directionality (for example from WC to CC), or breakpoints, were identified and quantified in two different ways: 1. Manually counting somatic breakpoints for each cell. 2. Using the computational tool BreakpointR, with a 2 Mb window-size parameter, to determine the genomic coordinates of the confidence interval for each breakpoint. The mouse genome was binned into 500 kb intervals, and breakpoint counts were quantified per bin. A cutoff of 3 breakpoints per bin in the *H2b*^fl/fl^ (WT) condition was applied, and bins meeting this criterion were classified as putative “germline” mutation regions. For a specific region on chromosome X (chrX:25,500,000-35,000,000), the boundaries were manually adjusted due to a highly segmented duplicated locus in the reference genome that inflated breakpoint counts. The resulting set of genomic regions was then used to filter breakpoints in *H2b*^fl/fl^, *H2b*^ΔIEC^ and *H2b*^ΔIEC^ /*STING*^-/-^ cells.

The aneuploidies were identified in the Strand-seq libraries by manually investigating the high-quality libraries. Specifically, changes in mean read depth were identified across large genomic regions, including whole-chromosome events, chromosome-arm-level alterations, and partial copy-number losses or gains exceeding 5 Mb. In Strand-seq data, aneuploidy can be inferred from strand-state imbalances by quantifying the Watson-to-Crick read ratio along the genome. Deviations from the expected ∼1:1 ratio indicate copy-number changes; for example, a Watson-specific gain is reflected by an increased Watson fraction, yielding an approximate Crick:Watson ratio of 1:2 across the affected region.

### RNA Sequencing (RNA-seq)

Total RNA was isolated from *H2b*^ΔIEC^ and *H2b*^ΔIEC^ /*STING*^-/-^ mouse organoids (n = 2 mice; three replicates per mouse) using the QIAGEN RNeasy Kit according to the manufacturer’s instructions. Samples were sequenced on HiSeq3000 (Illumina, San Diego, United States) using Illumina total RNA stranded TruSeq protocol. An average of ∼40 million 75-nt paired-end reads was sequenced for each sample. Raw reads were pre- processed using cutadapt (*46*) to remove adapter and low-quality sequences. RNAseq reads were aligned to the human genome (GrCh38) reference genome with TopHat2 (*47*). Gene expression values of the transcripts were computed by HTSe (*48*). Differential gene expression levels were analysed and visualized by the Bioconductor package DESeq2 (*49*). DeSeq2 was used to perform principal component analysis (PCA) that illustrate a separation according to the first principal component between different treatments. Venn diagrams were drawn using Venn Diagram package in R. Gene Ontology (GO) terms were obtained within the category of biological processes using the Innate DB database (www.innatedb.com) (*50*). Differential expression of genes between each treatment type versus control samples was determined using Wald tests to calculate the significance of differential expression for each gene in a pair-wise comparison between treatment types versus control.

### Statistics and reproducibility

Statistical analyses were performed using GraphPad Prism, and data are presented as mean ± SEM. Data distribution was first assessed using the Shapiro-Wilk normality test. For comparisons between two groups, an unpaired two-tailed Student’s t-test was applied when data followed a normal distribution; otherwise, the nonparametric Mann-Whitney U test was used. All statistical tests were two-sided, and p values < 0.05 were considered statistically significant. Additional statistical details are provided in the figure legends.

## Author Contributions

Conceptualization: KA, HX Methodology: KA, HX

Investigation: HX, MM, CW, NB, NAS, FW, JB, FT, NK, JB, SV, GY, QW, CP, JK, LW, BK, SS, PR, MT, KA

Supervision: PR, KA

Writing – original draft: HX, KA

Writing – review & editing: PR, SS, MT, AS, MM, NB

## Data, code, and materials availability

The data that support the findings of this study are available from the corresponding author upon reasonable request. Sequencing data will be deposited in a public repository and made available upon publication. Accession numbers will be provided prior to publication.

---

**Fig. S1.**
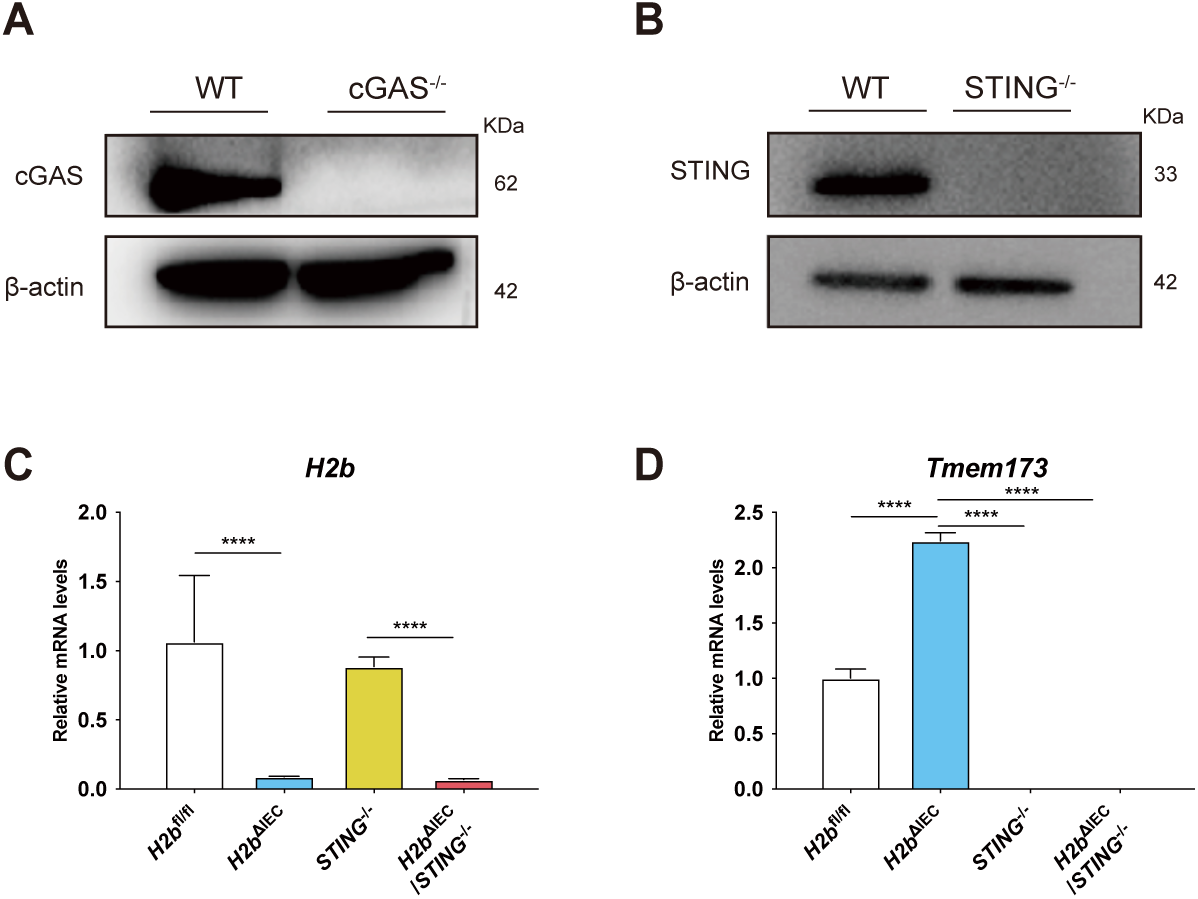
Genotype validation of knockout cell lines and mouse lines. **(A** to **B)** Western blot validation of MODE-K cGAS^-/-^ (A) and STING^-/-^ (B) knock out cell lines. **(C** to **D)** Genotyping of mouse lines using primers/probes targeting *H2b* (C) and *Tmem173* (D) in *H2b*^fl/fl^, *H2b*^ΔIEC^ ,*STING*^-/-^ and *H2b*^ΔIEC^/*STING*^-/-^ mice. *H2b* and *Tmem173* transcript levels across the four genotypes were quantified in parallel, as indicated. Significance was determined using unpaired two-tailed Student’s test. For all the significance analysis: ns = not significant, * p<0.05, ** p<0.01, *** p<0.001, **** p<0.0001.

**Fig. S2.**
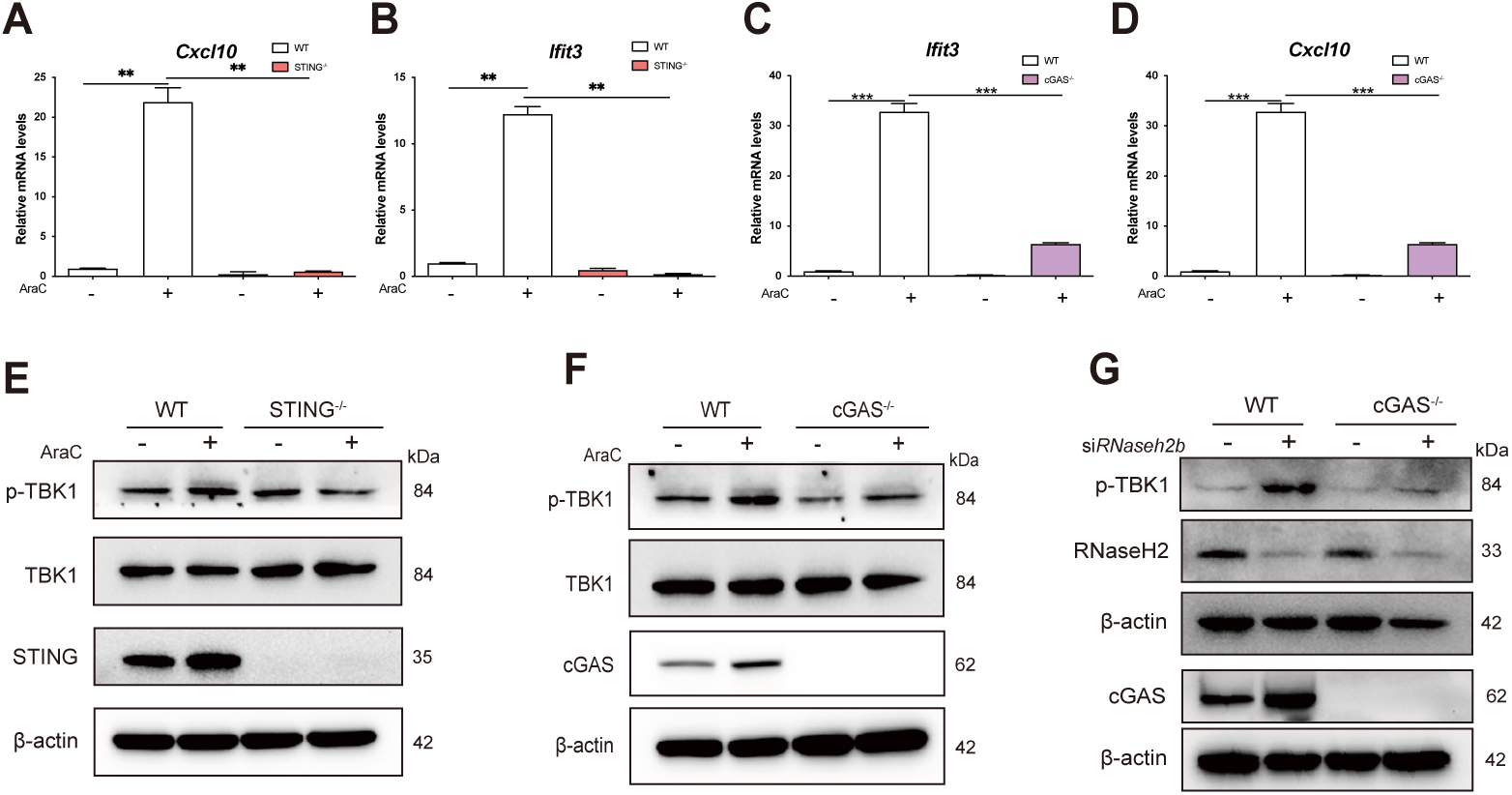
cGAS–STING-dependent IFN-I responses to DNA damage. **(A** and **D)** Gene expression of IFN-I pathway genes *Cxcl10* and *Ifit3* quantified by Taqman assay from MODE-K STING^-/-^ (A-B) or cGAS^-/-^ (C-D) cells were treated with 2.5 µM AraC for 20h, relative to MODE-K WT. Significance was determined using unpaired two-tailed Student’s test. **(E)** MODE-K WT and STING^-/-^ cells were treated with 2.5 µM AraC for 20h. Protein levels were measured by western blot assay. **(F)** MODE-K WT and cGAS^-/-^ cells were treated with 2.5 µM AraC for 20h. Protein levels were measured by western blot assay. **(G)** MODE-K WT and cGAS^-/-^ cells were transfected with either siRNA against *Rnaseh2b* or control siRNA for 24h. Protein levels were measured by western blot assay. For all the significance analysis: ns = not significant, * p<0.05, ** p<0.01, *** p<0.001, **** p<0.0001.

**Fig. S3.**
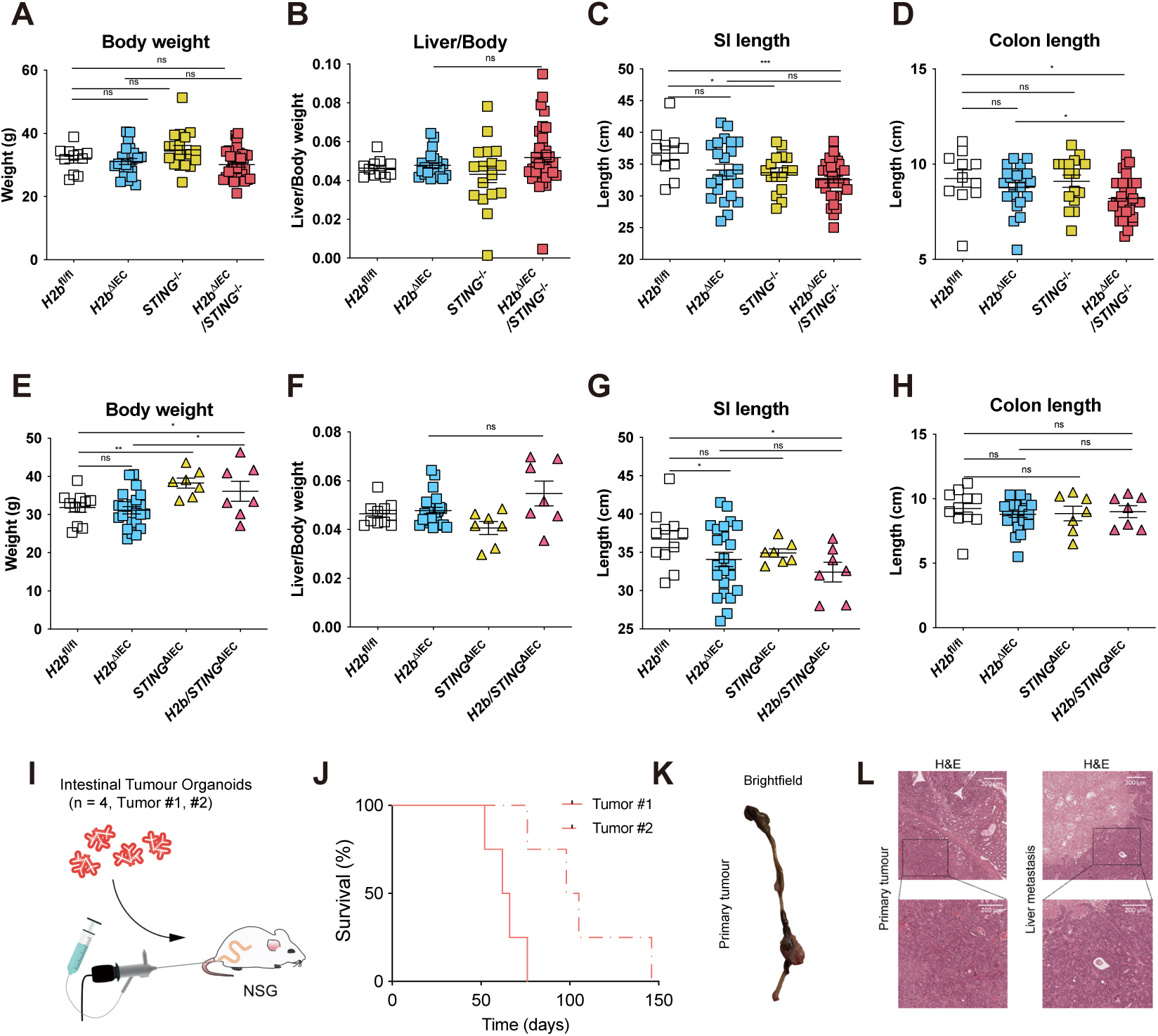
Clinical phenotype in aged *H2b*^ΔIEC^/*STING*^-/-^ mice. **(A** to **D)** Basal phenotyping analysis including Body weight (A), liver weight/body weight (B), Small intestine length (C), Colon length of aged (D) (> 53W) *H2b*^fl/fl^ (n=12), *H2b*^ΔIEC^ (n=23) ,*STING*^-/-^ (n=18) and *H2b*^ΔIEC^/*STING*^-/-^ (n=37) mouse line. Significance was determined using unpaired two-tailed Student’s test. **(E** to **H)** Clinical phenotype in aged *H2b*/*STING*^ΔIEC^ mice. Basal phenotyping analysis including body weight (E), liver weight/body weight (F), small intestine length (G), colon length (H) of aged (> 53W) *H2b*^fl/fl^ (n=12), *H2b*^ΔIEC^ (n=23) ,*STING*^ΔIEC^ (n=7) and *H2b*/*STING*^ΔIEC^ (n=7). *H2b*^fl/fl^ and *H2b*^ΔIEC^ mice are shared controls from the same cohort as shown in fig. S3 A-D. Significance was determined using unpaired two-tailed Student’s test. **(I)** Scheme of tumor organoid transplant model. **(J** to **L)** Survival curve of mice transplanted with tumor organoids isolated from *H2b*^ΔIEC^/*STING*^-/-^ mouse (J). Representative picture of primary tumor (K). H&E staining of primary and metastatic tumor (L). For the significance analysis: ns = not significant, * p<0.05, ** p<0.01, *** p<0.001, **** p<0.0001.

**Fig. S4.**
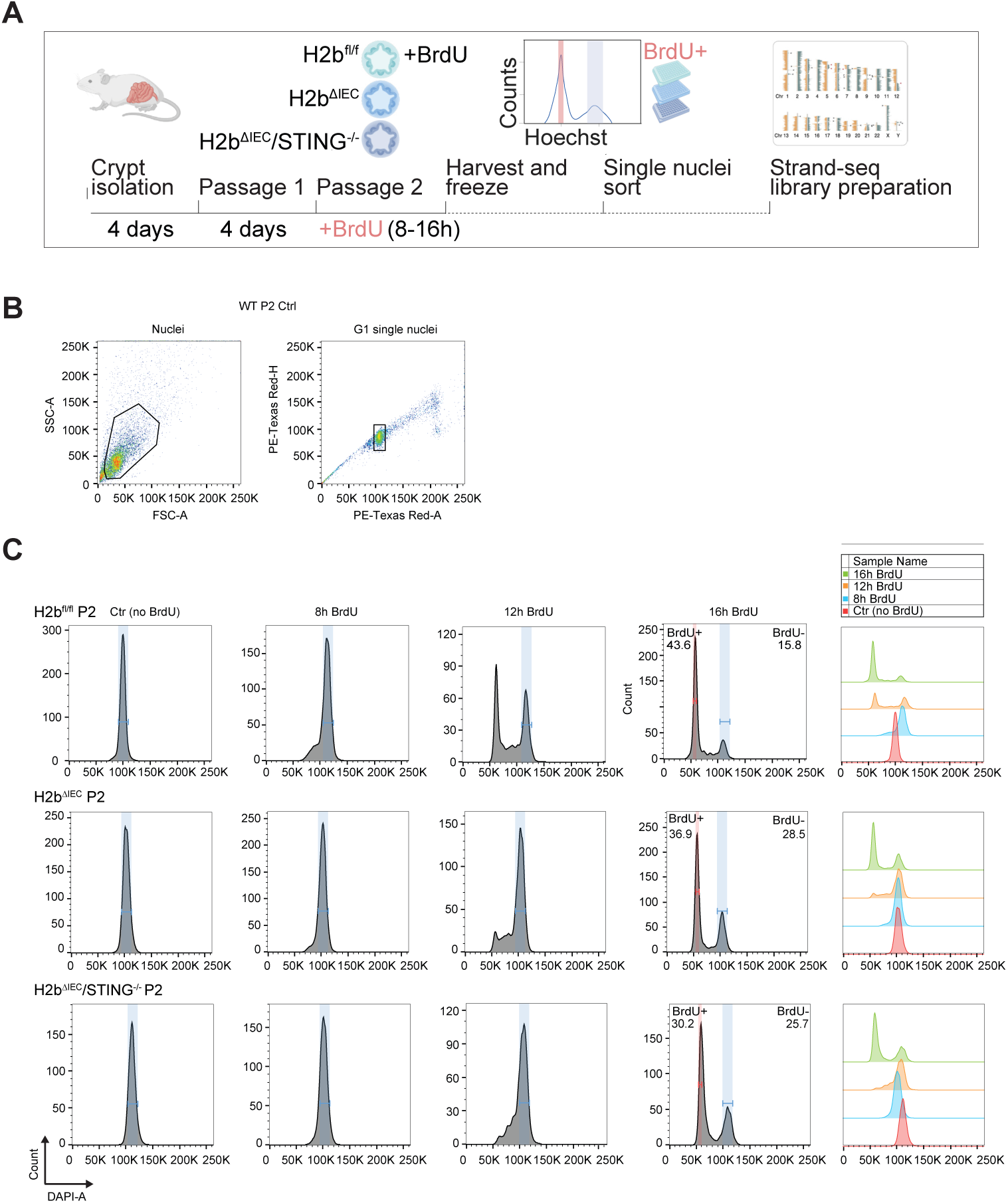
Experimental setup to produce Strand-seq libraries and FACS gating strategy. **(A)** Graphical design of experimental setup to produce Strand-seq libraries. **(B)** Example of the FACS gating used to select intact nuclei in G1 phase. **(C)** FACS gates showing the histogram counts of the DAPI-A channel for cells untreated with BrdU (Ctr), treated for 8, 12 or 16 hours with BrdU. The gate in blue shows the undivided peak (BrdU-), while the red gate indicates the nuclei of cells that divided once in presence of BrdU (hemi-substituted, BrdU+).

**Fig. S5.**
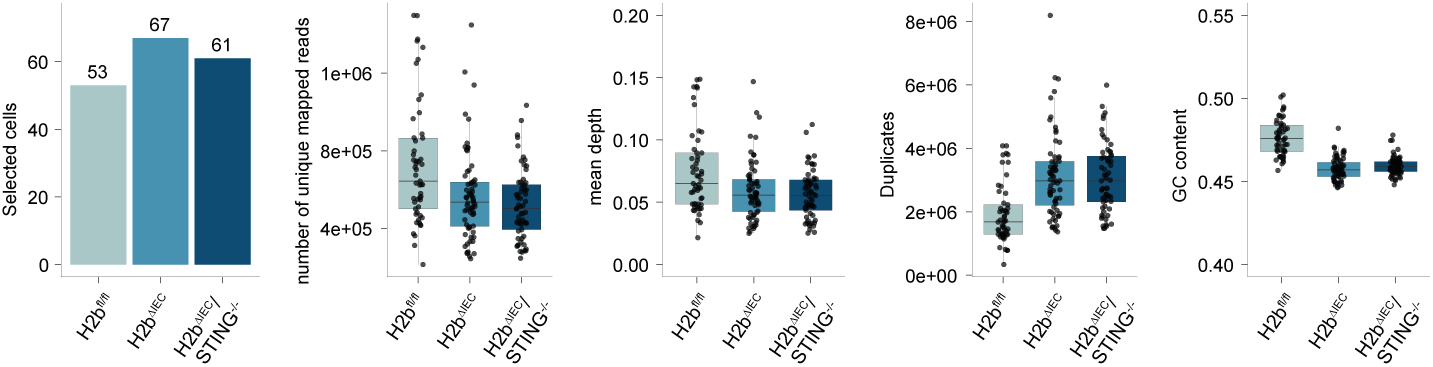
Strand-seq libraries quality. Histogram showing the number of selected cells per sample. The boxplots are showing the quality control of the selected libraries. Specifically showing the number of unique mapped reads, mean depth, duplicates and GC content. Each dot represents one single cell.

**Fig. S6.**
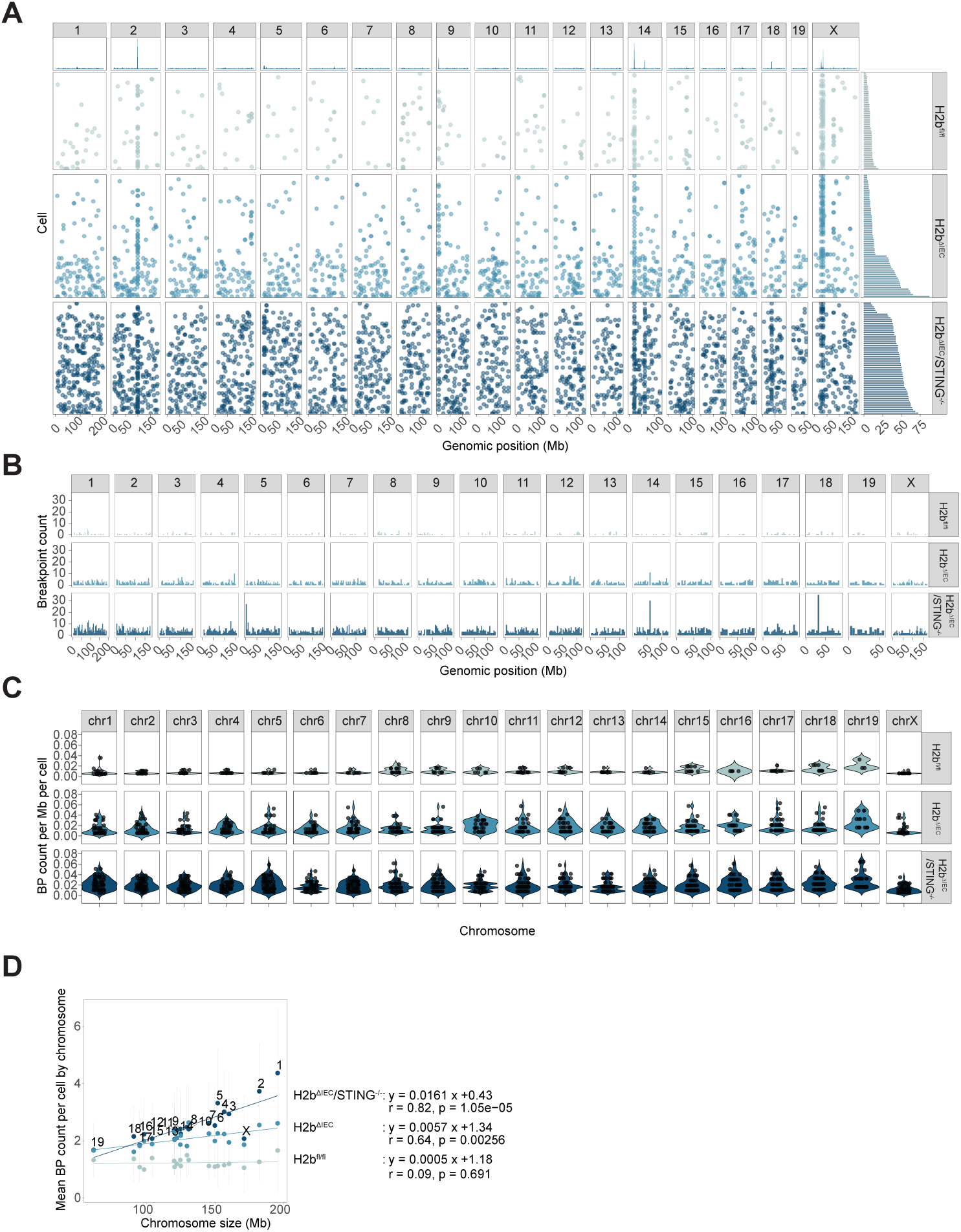
Breakpoint analysis and filtering strategy. **(A)** The plot is showing the breakpoints per cell by genomic location, identified with BreakpointR. Each dot represents a breakpoint. On the top panel the stack-histogram shows the quantification per genomic position (500 Kb bins). The panel on the right shows the histogram of the quantification of total breakpoint per cell. Chromosome Y was removed due to the low read mappability. **(B)** Histogram histogram shows the quantification of breakpoints per genomic position (4Mb binned genome) after filtering for *H2b*^fl/fl^ (WT) specific-recurrent regions. **(C)** Violin plot of the BP count per Mb by chromosome. **(D)** The plot shows the correlation between the chromosome size and the median BP count per chromosome. The grey bar represents the standard deviation for each chromosome and condition.

**Fig. S7.**
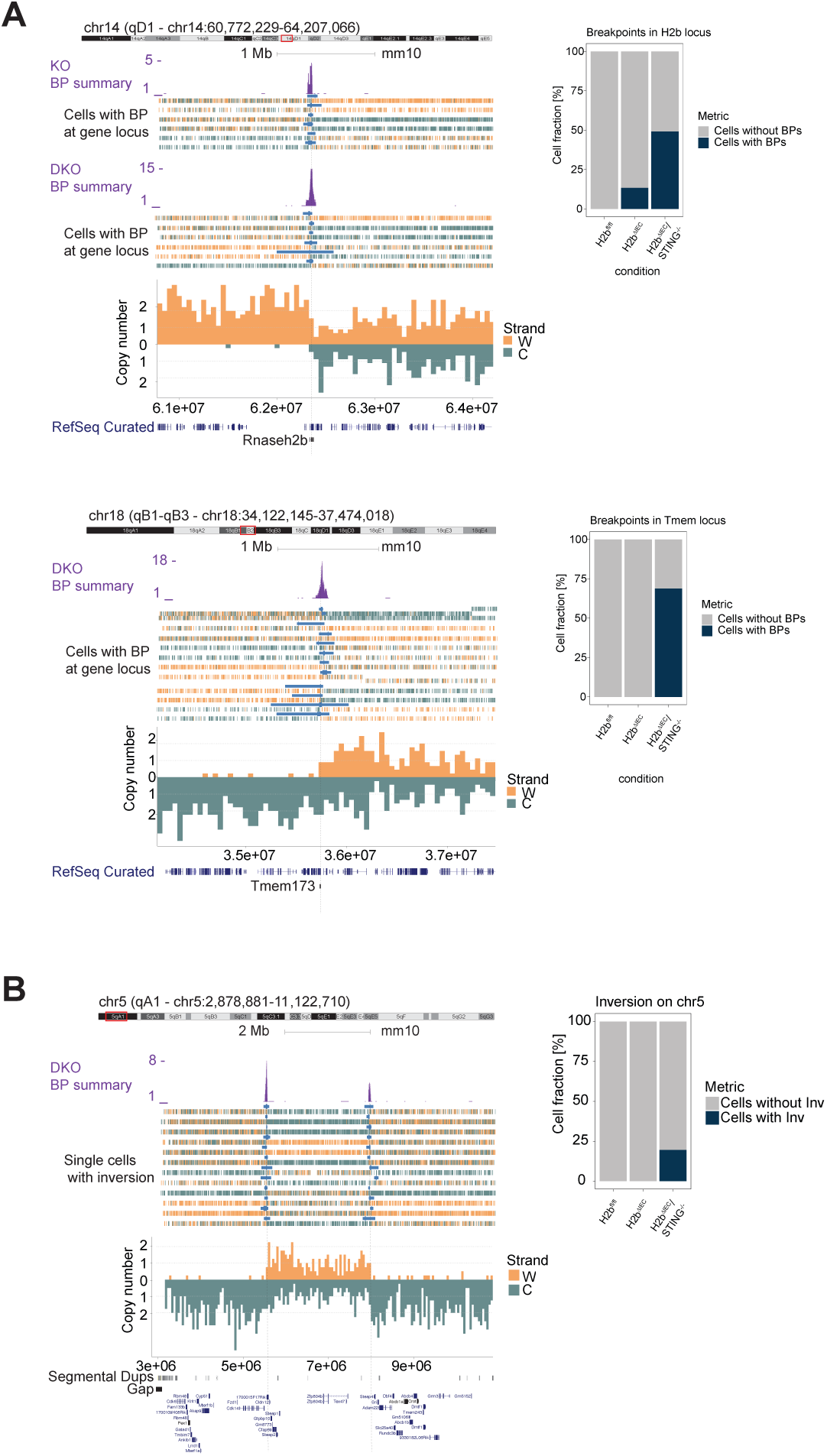
Breakpoint enrichment at specific loci. **(A)** UCSC visualization of the accumulation of breakpoints at *H2b* (left-upper panel) and *Sting* loci (left-lower panel). The violet histogram represents the overlapping BP count. The blue bar is the confidence interval of the BP per cell. Each row is a single cell, and the directionality can be visualized by the vertical bars that are binned reads (Watson= orange; Crick= green). At the bottom a mirrored plot of a representative cell possessing a BP at the gene locus. Right panels: Percentage of cell fraction with breakpoints on the H2b and Sting loci. **(B)** Representative ideograms of inverted region and UCSC visualization of the accumulation of breakpoints at the inversion borders. The violet histogram represents the overlapping BP count. The blue bar is the confidence interval of the BP per cell. Each row is a single cell, and the directionality can be visualized by the vertical bars that are binned reads (Watson= orange; Crick= green). Right panels: Percentage of cell fraction with inversion.

**Fig. S8.**
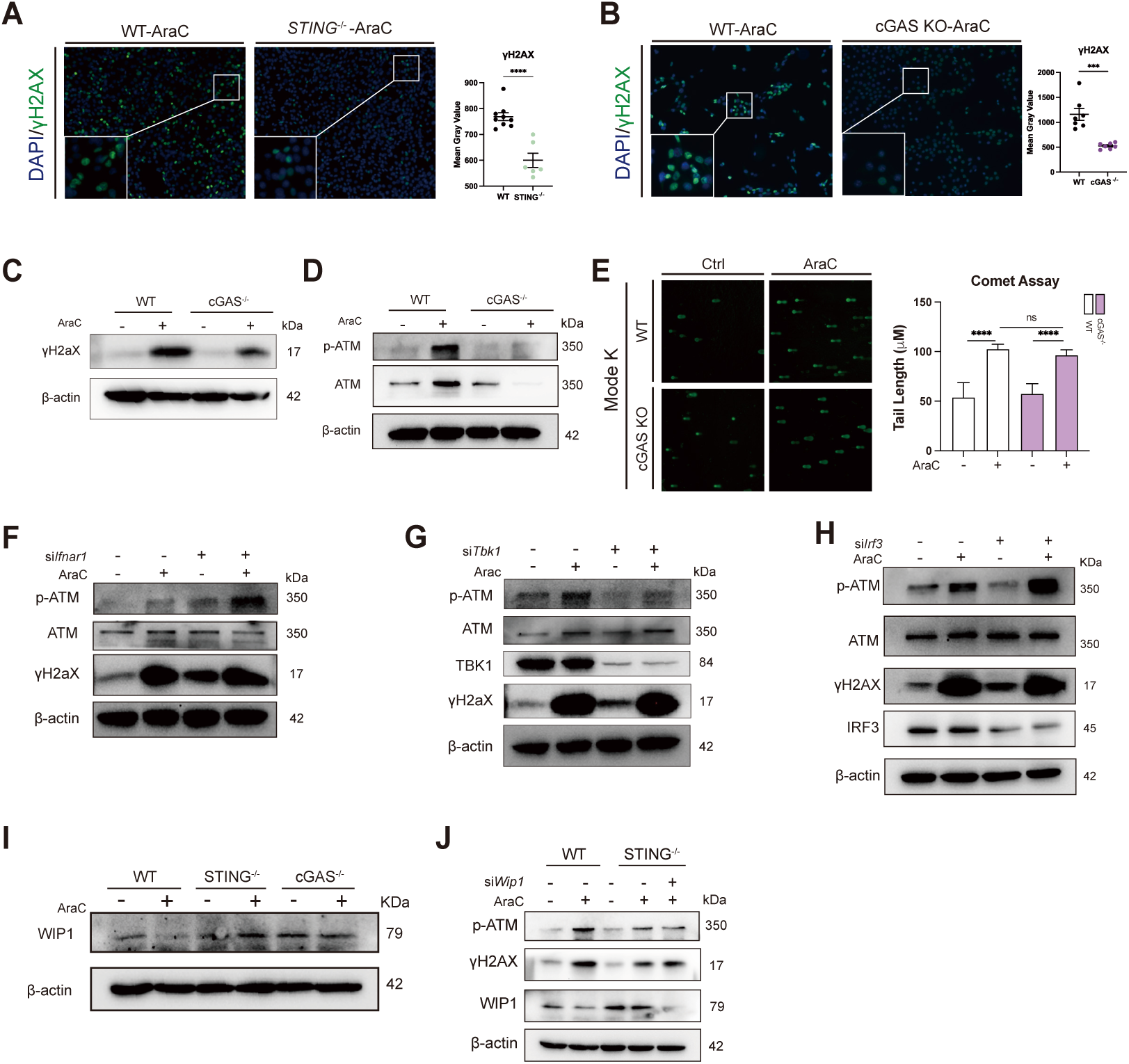
cGAS-STING pathway regulates HR in an IFN-I independent manner. **(A-B)** Immunofluorescence staining of γH2AX following AraC treatment in MODE-K cells. MODE-K STING^-/-^ (A) and MODE-K cGAS^-/-^ (B) cells were treated with 2.5 μM AraC for 20 h and stained for γH2AX by immunofluorescence. Representative images (Left) and quantification of nuclear γH2AX fluorescence intensity (Right) are shown for each genotype. Each dot represents the mean nuclear γH2AX fluorescence intensity across all cells within a single image. Significance was determined using unpaired two-tailed Student’s test. **(C-D)** HR related protein γH2AX (C) and ATM (D) assessed by western blot in MODE-K cGAS^-/-^ cells. **(E)** MODE-K cGAS^-/-^ cells were treated with 2.5 μM AraC for 20 h and subjected to a comet assay. DNA damage level was quantified by measuring comet tail length. Significance was determined using unpaired two-tailed Student’s test. **(F)** MODE-K WT, STING^-/-^, and cGAS^-/-^ cells were treated with 2.5 μM AraC and/or 100ng/ml IFN-β for 20 h, as indicated. Protein levels were analyzed by western blot. **(G** to **I**) **siRNA-mediated knockdown of canonical cGAS/STING pathway. MODE-K WT** cell was transfected with siRNAs targeting *Ifnar**1*****(G), *Tbk1*(H), *Irf3* (I)** or with a non-targeting control siRNA, then stimulated with 2.5 µM AraC for 20h. Protein levels were measured by western blot assay. **(J)** MODE-K WT, STING^-/-^, and cGAS^-/-^ cells were treated with 2.5 μM AraC for 20h. Protein levels were analyzed by western blot. For all the significance analysis: ns = not significant, * p<0.05, ** p<0.01, *** p<0.001, **** p<0.0001.

**Fig. S9.**
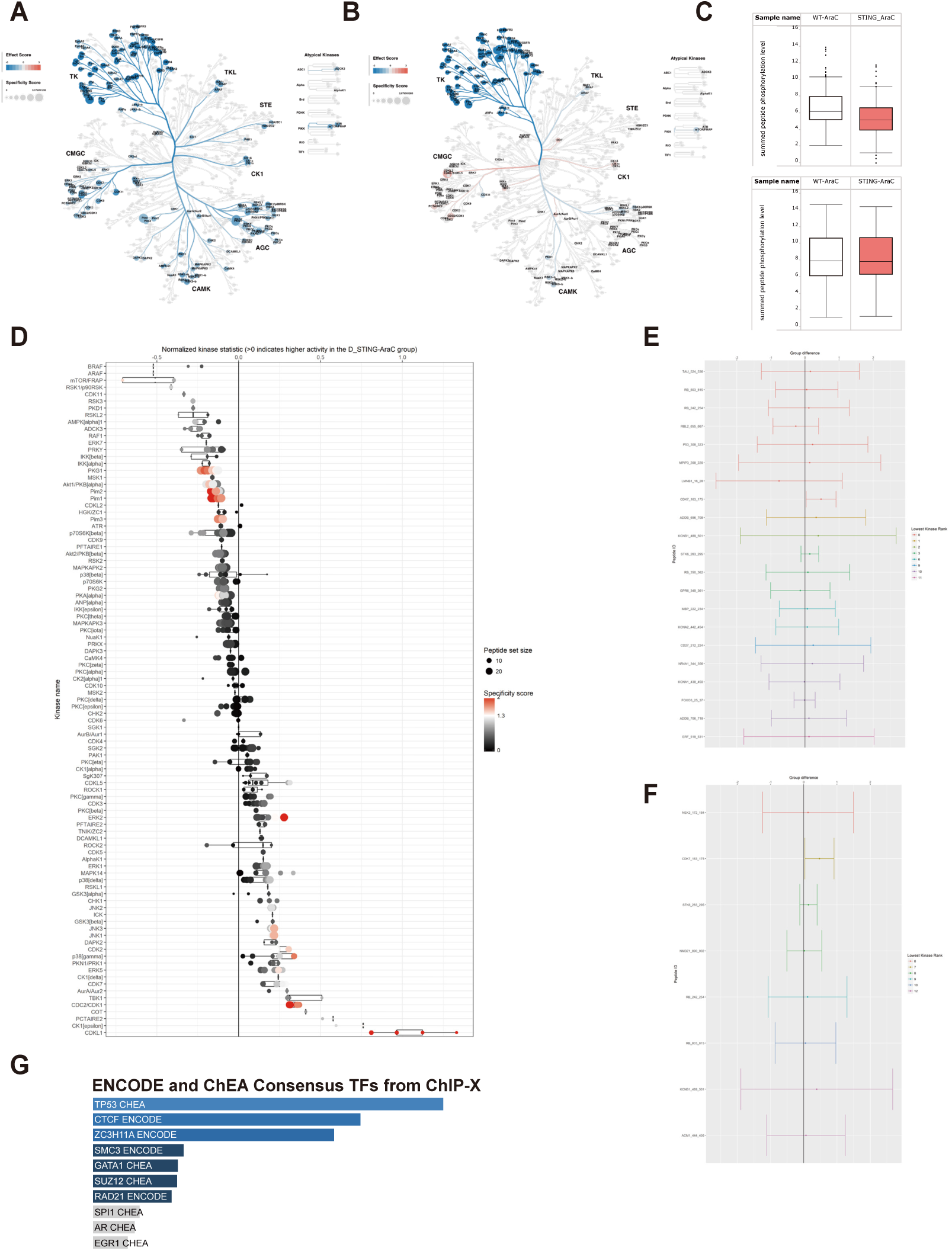
Phospho-kinase profiling reveals altered kinase activity networks in MODE-K STING^-/-^ cells under control and AraC treatment. **(A** and **B)** Kinome tree visualization of upstream kinase activity in MODE-K cells. Upstream kinase analysis results are mapped onto the kinome tree, comparing MODE-K STING^-/-^ with WT under control (A) conditions and AraC (B) treatment. Colors indicate relative kinase activity in MODE-K STING^-/-^ versus WT (blue, decreased; red, increased). **(C)** Box plots showing the sum of log2-transformed fluorescence signal intensities of tyrosine kinase (PTK) phosphorylation levels on the chip. **(D)** Upstream kinase analysis based on degree of phosphorylation from serine-threonine kinase assay (STK) of MODE-K STING^-/-^ compared with WT. Based on activity and phosphorylation pattern a prediction of the active kinase is given by the BioNavigator 6 software. Size of the peptide set is shown by dot size. Color code ranges from red (high specificity) to black (low specificity) and kinases are sorted by specificity score. **(E** and **F)** Representative peptides supporting kinase assignment. Representative peptide signals are shown for (E) CDK1 and (F) CDKL1. **(G)** EnrichR pathway enrichment based on upstream kinase analysis. Kinases inferred to have decreased or increased activity in AraC treated MODE-K STING^-/-^ compared with WT were converted to gene names and used as input for EnrichR. Bar plots show enriched terms ranked by significance (logP; top to bottom), with decreased-activity kinases shown in the upper panel and increased-activity kinases shown in the lower panel.

**Fig. S10.**
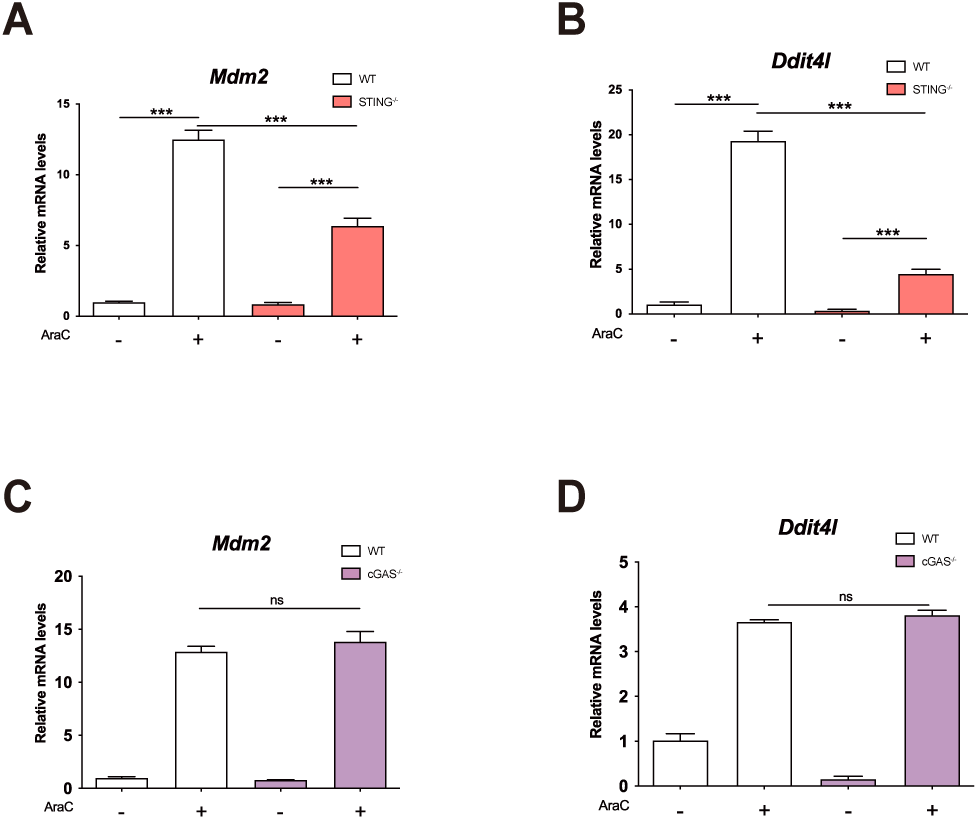
Differential roles of STING and cGAS in regulating DNA damage-induced p53 signaling via *Ddit4l*. **(A** and **B)** Gene expression of *Mdm2* and *Ddit4l* quantified by Taqman assay from MODE-K STING^-/-^ cells treated with 2.5 µM AraC for 24h, relative to MODE-K WT. Significance was determined using unpaired two-tailed Student’s test. **(C** and **D)** Gene expression of *Mdm2* and *Ddit4l* quantified by Taqman assay from MODE-K cGAS^-/-^ cells treated with 2.5 µM AraC for 24h, relative to MODE-K WT. Significance was determined using unpaired two-tailed Student’s test. For all the significance analysis: ns = not significant, * p<0.05, ** p<0.01, *** p<0.001, **** p<0.0001.

**Fig. S11.**
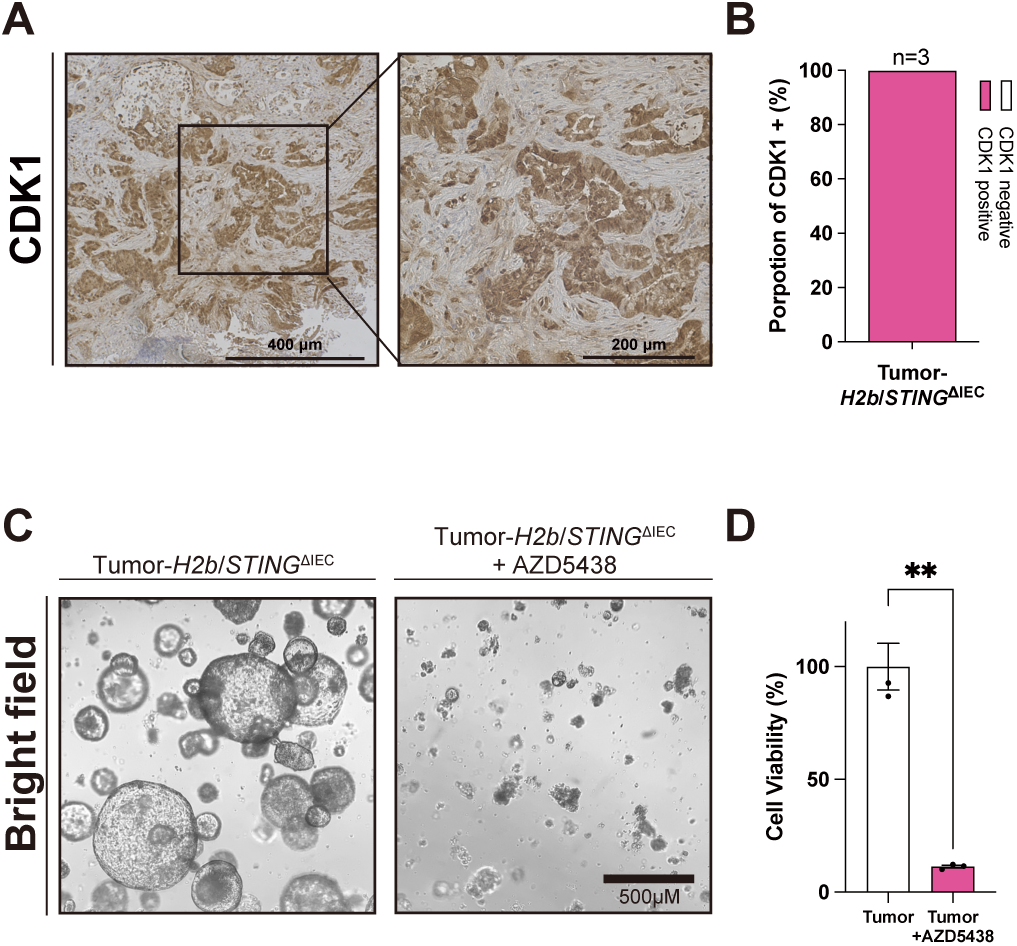
STING-deficient tumors exhibit elevated CDK1 activity and vulnerability to CDK inhibition. **(A** and **B)** Immunohistochemical staining of CDK1 in tumor tissues derived from H2b/STING^ΔIEC^ mice. Representative picture of CDK1 staining (A). Proportion of CDK1-positive tumors (B). **(C** and **D)** Representative pictures (C) and CellTiter-Glo assay (D) from *H2b*/*STING* ^ΔIEC^ tumor organoids treated with Control (DMSO) or 1 μM AZD5438 for 72h. Significance was determined using unpaired two-tailed Student’s test. For all the significance analysis: ns = not significant, * p<0.05, ** p<0.01, *** p<0.001, **** p<0.0001.

**Fig. S12.**
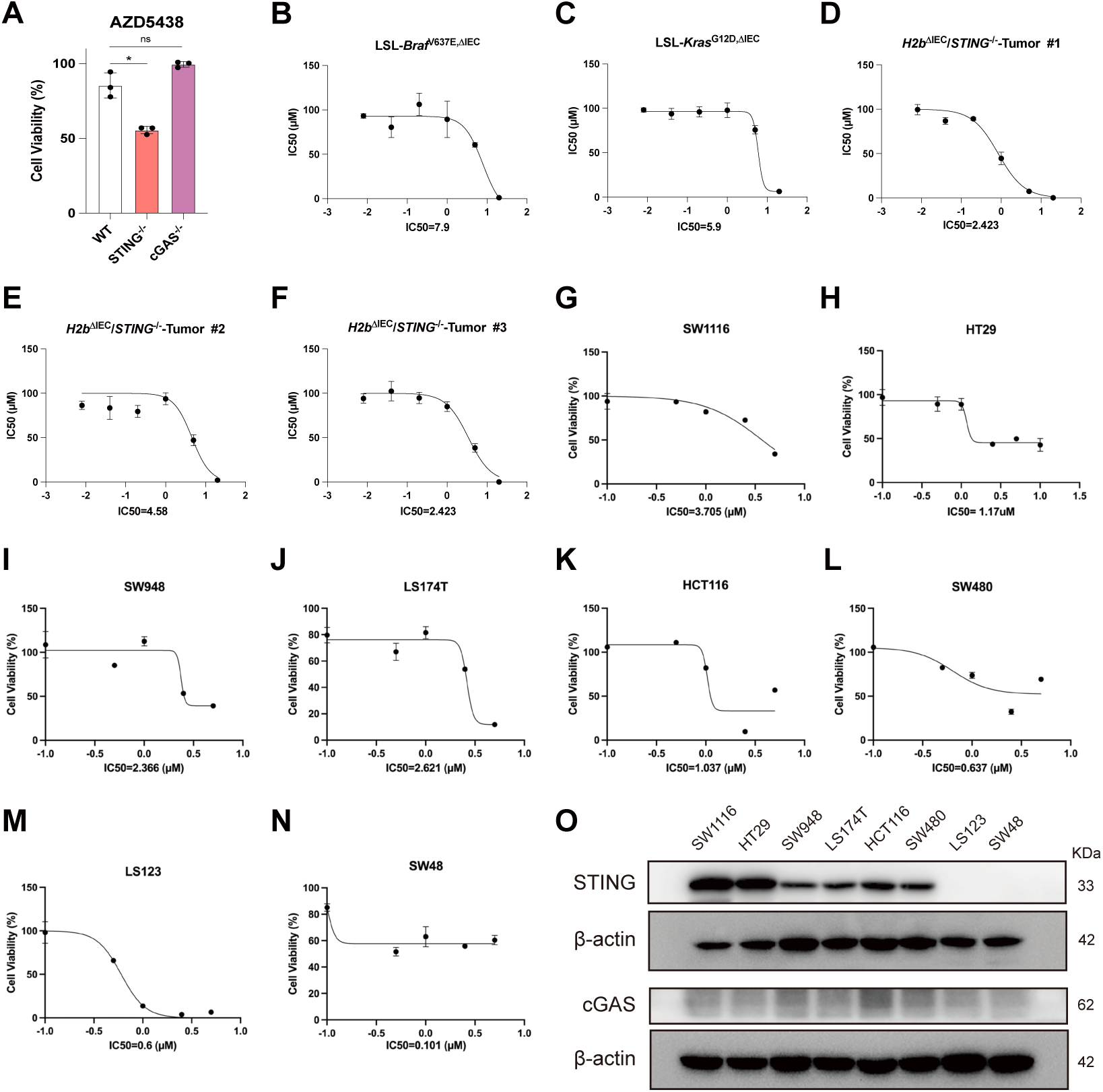
STING axis status associates with CDK inhibitor response in mouse tumor organoids and human CRC cell lines. **(A)** Cell viability following AZD5438 treatment in MODE-K cell lines. MODE-K WT, STING-/-, and cGAS-/- cells were treated with 1 μM AZD5438 for 72 h. Cell viability was assessed using the CellTiter-Glo luminescent cell viability assay. Significance was determined using unpaired two-tailed Student’s test. **(B** to **F)** AZD5438 dose–response and IC50 determination in tumor organoids. Tumor organoids were treated with a gradient of AZD5438 (Ctrl, 0.25, 0.5, 1, 2.5, and 5 μM) for 3 days. Cell viability was quantified using the CellTiter-Glo luminescent assay, and dose–response curves were used to calculate IC50 values. IC50 values in SI tumor organoids derived from LSL-Braf^V637E,ΔIEC^ mouse line (B). IC50 in LSL-Kras^G12D,ΔIEC^ tumor organoids (C). IC50 values in different *H2b*/*STING*^ΔIEC^ mouse (D-F). **(G** to **N)** AZD5438 dose–response and IC50 determination in human colon adenocarcinoma cell lines. Indicated human colon adenocarcinoma cell lines were treated with a gradient of AZD5438 (Ctrl, 0.25, 0.5, 1, 2.5, and 5 μM) for 3 days. Cell viability was quantified using the CellTiter-Glo luminescent assay, and dose–response curves were generated to calculate IC50 values for each cell line. **(O)** cGAS-STING protein levels in human colon adenocarcinoma cell lines were assessed by western blot. For all the significance analysis: ns = not significant, * p<0.05, ** p<0.01, *** p<0.001, **** p<0.0001.

**Table S1:** Antibody List.

| **Antibody** | **Source** | **Catalog** | **Dilution** | **Usage** |
| --- | --- | --- | --- | --- |
| STING | Cell Signaling Technology | 13647S | 1:1000 | WB |
| ATM | Cell Signaling Technology | 2873T | 1:1000 | WB |
| p-ATM | santacruz | sc-47739 | 1:1000 | WB |
| cGAS | Cell Signaling Technology | 31659S | 1:1000 | WB |
| P53 | Cell Signaling Technology | 2524S | 1:1000 | WB |
| TBK1 | Cell Signaling Technology | 3013S | 1:1000 | WB |
| p-TBK1 | Cell Signaling Technology | 5483S | 1:1000 | WB |
| γH2AX | Cell Signaling Technology | 2577S | 1:1000 | WB |
| WIP1 | Cell Signaling Technology | 11901 | 1:1000 | WB |
| cdc2 (CDK1) | Cell Signaling Technology | 28439S | 1:1000 | WB |
| p-cdc2(p-CDK1 T161) | Cell Signaling Technology | 9114S | 1:1000 | WB |
| P21 | Cell Signaling Technology | 37543S | 1:1000 | WB |
| RnaseH2 | Shared by collaborator (2) | - | 1:1000 | WB |
| NBS1 | Cell Signaling Technology | 14956 | 1:1000 | WB |
| β-actin | Cell Signaling Technology | 4970S | 1:3000 | WB |
| IRF3 | abcam | ab76493 | 1:1000 | WB |
| mouse HRP | Amersham Biosciences | NA931V | 1:2000 | WB |
| rabbit HRP | Amersham Biosciences | NA934V | 1:2000 | WB |
| p-ATM | santacruz | sc-47739 | 1:100 | IF |
| γH2AX | Cell Signaling Technology | 2577S | 1:500 | IF |
| Alexa Fluor 488-goat anti-rabbit | Thermo Fisher Scientific | A11070 | 1:500 | IF |
| cdc2 (CDK1) | Cell Signaling Technology | 28439S | 1:500 | IHC |
| rabbit HRP | Amersham Biosciences | NA934V | 1:2000 | IHC |
| NBS1 | Cell Signaling Technology | 14956 | 1:100 | co-IP |
| ATM | abcam | ab201022 | 1:100 | co-IP |

**Table S2.** Primer list.

| **Gene name** | **Source** | **Ref ID** |
| --- | --- | --- |
| *Rnaseh2b* | IDT | Mm.PT.56a.41665482 |
| *β-actin (Actb)* | IDT | Mm.PT.39a.22214843.g |
| *Tmem173* | IDT | Mm.PT.58.12798185 |
| *Mdm2* | IDT | Mm.PT.58.42166864 |
| *Ddit4l* | IDT | Mm.PT.58.32152752 |
| *Cxcl10* | IDT | Mm.PT.58.43575827 |
| *Ifit3* | IDT | Mm.PT.58.33537107 |

**Table S3.** siRNA list.

| **Gene name** | **Source** | **Catalog** |
| --- | --- | --- |
| *Rnaseh2b* | QIAGEN | GS67153 |
| *Ddit4l* | QIAGEN | GS73284 |
| *Tmem173* | QIAGEN | GS72512 |
| *cgas* | QIAGEN | GS214763 |
| *Wip1* | Thermofisher | s79351 |
| *Ifnal1* | QIAGEN | GS15975 |
| *Irf3* | QIAGEN | GS54131 |
| *Trp53* | QIAGEN | GS22059 |

**Table S4.** stimulants list.

| Name | Source | Catalog |
| --- | --- | --- |
| AraC | Sigma-Aldrich | C6645-25mg |
| AZD5438 | MCE | HY-10012 |
| AZD0156 | MCE | HY-100016 |
| rm IFN-beta Protein (with carrier) | R&D Systems | 8234-MB-010 |

